# Filopodome proteomics identifies CCT8 as a MYO10 interactor critical for filopodia functions

**DOI:** 10.64898/2025.12.03.691809

**Authors:** Ana Popović, Neil J. Ball, Mitro Miihkinen, Omkar Joshi, Michal Dibus, Marjaana Ojalill, Joanna Pylvänäinen, Johanna Ivaska, Benjamin T. Goult, Guillaume Jacquemet

## Abstract

Cancer cells utilize filopodia to explore, adhere to, and invade their surrounding microenvironment, yet the protein networks that organize these protrusions remain incompletely defined. To uncover the molecular machinery underlying MYO10-positive filopodia, we targeted the fast biotin ligase TurboID to the filopodia tip-localized motor protein MYO10. Proximity biotinylation in two cell types revealed hundreds of potential MYO10 interactors. Surprisingly, there was limited overlap between the cell lines, indicating a previously unknown level of cell-type specificity in filopodia composition. A targeted microscopy and siRNA screen identified MINK1, SCRIB, CSNK1A1, and CCT8 as new regulators of filopodia formation. Focusing on one common interactor between cell lines, CCT8, known as a subunit of the chaperonin TRiC (TCP1 Ring Complex), we found that CCT8 associates with the MYO10 motor domain and regulates MYO10 filopodia independently of TRiC. Depleting CCT8 affected filopodia dynamics and impaired cell spreading, migration, and invasion in breast cancer cells. These findings establish CCT8 as a TRiC-independent regulator of MYO10 filopodia across different cancer cell types, highlight the surprising cell-type-specificity of filopodia composition, and provide a strategy and resource for studying filopodia in various biological contexts.

## Introduction

Filopodia are widespread, small, dynamic, actin-rich projections, usually measuring 1-5 µm in length and 50-200 nm in diameter, that serve as the initial contact point between a cell and its environment (Blake and Gallop, 2023). Filopodia contain receptors such as integrins, cadherins, and growth factor receptors, which detect biochemical and mechanical signals from the extracellular matrix (ECM) and neighboring cells. By sensing the external cellular environment, filopodia play essential roles in in vitro and in vivo processes, including development, angiogenesis, and immune surveillance (Jacquemet et al., 2015).

Despite their importance, the molecular composition of filopodia remains mostly unknown. We and others have used targeted microscopy screens to identify specific proteins and lipid species at filopodia tips (Jacquemet et al., 2019; Blake et al., 2024; Sudhaharan et al., 2019; Mason et al., 2025) and, for example, have identified specific adhesion molecules that accumulate at filopodia (Jacquemet et al., 2015, 2016). However, microscopy methods are inherently biased toward known or expected proteins and cannot uncover the entire “filopodome.” In contrast, other cell adhesion and signaling hubs, such as focal adhesions, have been successfully isolated and identified by using mass spectrometry-based techniques, including proximity biotinylation strategies (Chastney et al., 2020; Atherton et al., 2022; Dong et al., 2016), leading to comprehensive “adhesome” datasets that have advanced the field. Unbiased characterisation of protrusions, such as invadopodia, has also been successfully performed using proximity biotinylation strategies (Thuault et al., 2020). Similar proteomic analyses of filopodia are lacking, mainly because of their small size and transient nature, which make biochemical isolation difficult. As a result, filopodia are often viewed as uniform structures, even though different filopodia populations can contain unique protein groups and show varied dynamics and functions (Jacquemet et al., 2019; Young et al., 2018).

Filopodia assembly is driven by linear actin polymerization, with barbed ends oriented toward the plasma membrane, and is bundled by actin crosslinkers. This structure allows unconventional myosins, such as MYO10, to move processively toward and accumulate at filopodia tips (Zhang et al., 2004). In addition to MYO10 physiological roles, MYO10 has been connected to pathological conditions. For example, MYO10 expression is elevated in several cancers, where it supports cancer cell migration, invasion, and metastasis (Arjonen et al., 2014). Although some MYO10 interactors are known (Zhang et al., 2004; Popović et al., 2023; Wei et al., 2011; Bennett and Strehler, 2008), a complete interactome for MYO10 remains unavailable.

Here, to address these gaps, we employed TurboID-mediated proximity biotinylation fused to MYO10, leveraging its strong localization at filopodia tips as a molecular handle to label proteins at these sites. Applying this method to two biologically and biomechanically distinct cancer cell lines (osteosarcoma U-2 OS and glioblastoma U-87 MG), we identified hundreds of candidate proteins with limited overlap between the cell lines, underscoring the cell-type-dependent organization of MYO10 filopodia. Using targeted microscopy and an siRNA screen, we validated MINK1, SCRIB, CSNK1A1, and CCT8 as new regulators of filopodia formation. We focused on CCT8, a subunit of the TRiC (TCP1 Ring Complex) (Zeng et al., 2024), and showed that it regulates MYO10 filopodia independently of TRiC functionality by directly binding to the MYO10 motor domain. Phenotypically, CCT8 depletion decreased filopodia stability and impaired cell spreading, migration, and invasion in breast cancer cells. Overall, our study presents a robust strategy for profiling the molecular composition of MYO10 filopodia, offers a dataset for future research, and uncovers an unexpected TRiC-independent role for CCT8 in controlling filopodia.

## Results

### Profiling of MYO10 filopodia and MYO10 interactome by TurboID reveals cell-type-specific networks

To systematically characterize the composition of MYO10-positive filopodia and identify new MYO10 interactors, we used a proximity-dependent biotinylation method. Previously, we used BioID fused to MYO10 to isolate filopodia-related proteins (MYO10-BioID dataset); however, this method identified only a few proteins (Popović et al., 2023), likely due to BioID’s slow biotinylation kinetics. Since filopodia are highly dynamic, we chose TurboID here, as it enables faster biotinylation and may be better suited for capturing transient protein interactions (Branon et al., 2018).

We created a TurboID-MYO10 construct by fusing TurboID to the N-terminus of human MYO10 (Fig. 1A) and expressed TurboID-MYO10 in two different cell models: U-2 OS (osteosarcoma) and U-87 MG (glioblastoma). These cell lines effectively expressed the TurboID-MYO10 construct, allowing for the generation of stable cell lines suitable for large-scale mass spectrometry analysis. In cells expressing TurboID-MYO10, we observed clear and specific biotinylation at filopodia tips after a 1-hour biotin treatment, confirming the proper localization and enzymatic activity of our construct (Fig. 1B). Subsequently, biotinylated proteins were isolated using streptavidin pull-down assays from both cell lines treated with biotin or avidin (negative control), then proteins were separated by SDS-PAGE, visualized by western blotting (Fig. 1C and Fig. S1A) and identified by mass spectrometry (TurboID-MYO10 datasets, Fig. 1D, Supplementary table 1). Using this strategy, we identified proteins previously known to interact directly or indirectly with MYO10 at filopodia tips, including RAPH1 (also known as lamellipodin; (Popović et al., 2023)), ITGB1 (Integrin β1; (Zhang et al., 2004)), and TLN1 (Talin-1; (Miihkinen et al., 2021)). Additionally, proteins such as FLNA (Filamin-A) and AHNAK were also identified here, as in our previous MYO10-BioID dataset (Popović et al., 2023).

**Fig. 1.**
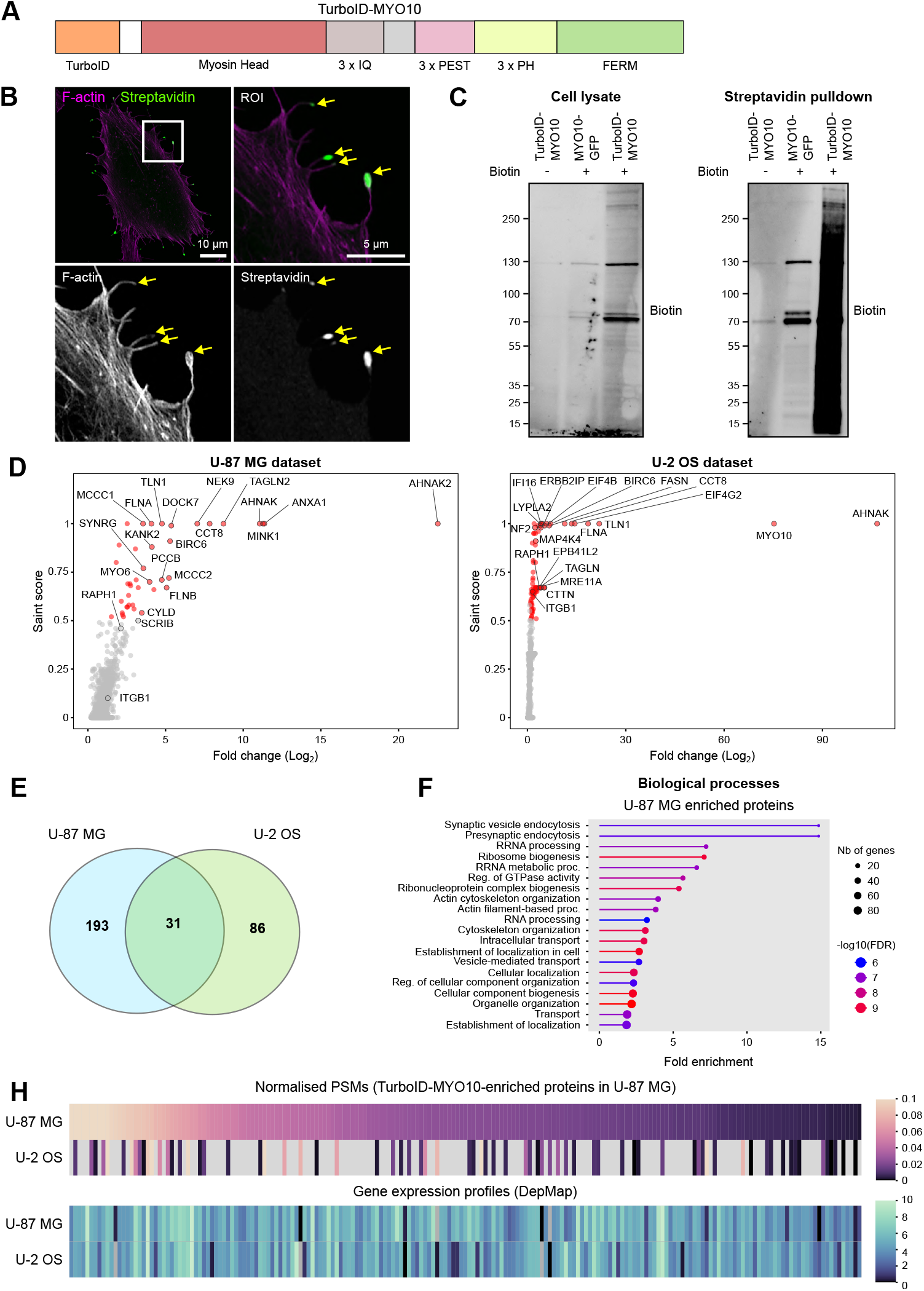
Identification of MYO10-associated proteins using proximity-dependent biotinylation. (**A**) Schematic representation of the TurboID-MYO10 construct used, highlighting TurboID fused to the N-terminus of MYO10. (**B**) U-87 MG cells stably expressing TurboID-MYO10 were seeded onto fibronectin-coated coverslips, and biotinylation of proximal proteins was induced by adding exogenous biotin for 1 hour. Cells were fixed 2 hours post-seeding (1-hour post-biotin addition), stained using fluorescent streptavidin (to detect biotinylated proteins) and phalloidin (to visualize F-actin), and imaged by confocal microscopy (Zeiss LSM880). Scale bars: Main, 10 µm; ROI, 5 µm. (**C–F**) U-87 MG and U-2 OS cells stably expressing TurboID-MYO10 were cultured on fibronectin-coated dishes for 2 hours, with biotinylation initiated 1 hour before lysis. Biotinylated proteins were affinity-purified using streptavidin-coated beads and analyzed by western blotting and mass spectrometry. (**C**) Representative western blot from U-87 MG cells displaying biotinylated protein profiles. (**D**) Volcano plots depicting proteins significantly enriched in proximity to MYO10 in U-87 MG and U-2 OS cells, respectively. The plots illustrate the significance (SAINT score calculated via CRAPome) on the x-axis versus fold enrichment relative to control on the y-axis. (**E**) Venn diagram summarizing the overlap and cell line-specific proteins enriched in U-87 MG and U-2 OS datasets. (**F**) Gene ontology analysis highlighting biological processes significantly enriched among proteins identified in U-87 MG cells. (**G**) Heatmaps comparing detected MYO10-associated proteins across the two cell lines, highlighting those proteins enriched in the U-87 MG dataset. Expected expression profiles derived from the DepMap database are included for comparison. The numerical data and the raw images used to make this figure have been archived on Zenodo (https://zenodo.org/records/17779715).

Using an enrichment cut-off criterion based on prior knowledge of known MYO10 interactors (see method for details), we identified 224 proteins in U-87 MG cells and 117 proteins in U-2 OS cells as potential MYO10 binders (Fig. 1E). Besides canonical cytoskeletal and adhesion components (FLNA, TLN1/2, TAGLN/TAGLN2, MYH9/14, NF2, DLG5, EPB41L1/2, TJP1/2, and SCRIB), the proximity labeling datasets highlight an extensive network of proteins involved in controlling membrane composition and lipid signaling. These include multiple phosphoinositide-metabolizing enzymes (PIP4K2A/C, PIP5K1C, MTMR2, SYNJ1/2), oxysterol-binding proteins (OSBPL6, OSBPL7, OSBPL8), enzymes involved in fatty acid and glycerophospholipid pathways (FASN, LYPLA2, CEPT1, PGAP1), and several lipid-binding adaptors and scaffolds (ANXA1/2, FABP5, PHLDB1, PLEKHF1, LANCL2). Surprisingly, only 31 proteins overlapped between the TurboID-MYO10 datasets, indicating unexpected cell-type specificity in potential MYO10 interactors (Fig. 1E). Gene ontology analysis further revealed that the enriched proteins in each cell line participate in different biological processes (Fig. 1F and Fig. S1B).

We then investigated whether differences in gene expression could explain the distinct sets of protein interactors observed in the U-87 MG and U-2 OS TurboID-MYO10 datasets. To do this, we combined our mass spectrometry data with mRNA expression profiles from U-87 MG and U-2 OS cells (DepMap database, (Arafeh et al., 2025)). Remarkably, side-by-side heatmap comparisons of TurboID-MYO10 protein levels and corresponding mRNA levels (Fig. 1G and Fig. S1C) showed that transcript expression alone does not account for the cell line–specific interactomes. Instead, these findings suggest additional regulatory layers, such as variations in interaction network architecture or post-translational modifications, influence the recruitment of proteins to MYO10-positive filopodia.

### MINK1, SCRIB, CSNK1A1, and CCT8 modulate filopodia formation

To demonstrate how the TurboID-MYO10 datasets can be mined to identify novel filopodia regulators, we chose six candidate proteins: CCT8 and SCRIB (enriched in both U-2 OS and U-87 MG cells), NF2, MAP4K4, CSNK1A1 (enriched in U-2 OS cells), and MINK1 (enriched in U-87 MG cells) for further investigation. These proteins were selected based on their enrichment profiles and because of known links to the actin cytoskeleton.

First, we determined the subcellular localization of these candidate proteins. Each protein was tagged with GFP and coexpressed with mScarlet-MYO10 in U-87 MG cells. Using structured illumination microscopy and Airyscan confocal microscopy, we examined the localization patterns of these proteins (Fig. 2A). All six candidate proteins localized to filopodia. Notably, MAP4K4, MINK1, SCRIB, and CSNK1A1 showed distinct enrichment at filopodia tips. In contrast, CCT8 and NF2 showed a more diffuse distribution across both filopodia and the cytoplasmic compartments.

**Fig. 2.**
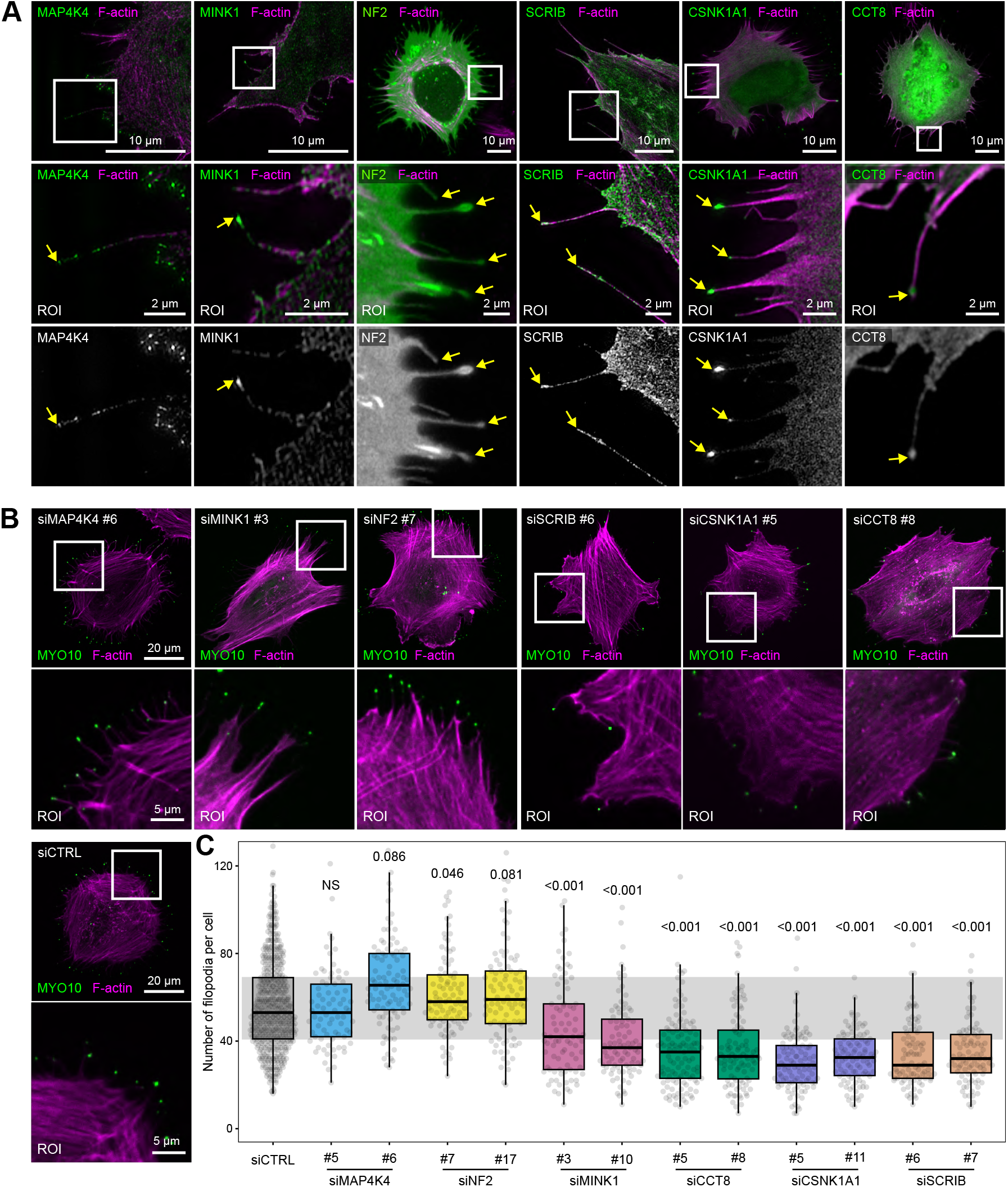
Localization and functional screening of filopodia-associated proteins identified by TurboID-MYO10. (**A**) Representative images of U-87 MG cells expressing MYO10-mScarlet and the indicated GFP-tagged candidate proteins (MAP4K4, MINK1, NF2, SCRIB, CSNK1A1, or CCT8). Cells were plated on fibronectin-coated coverslips for 2 hours, then fixed and imaged using either structured illumination microscopy (SIM) or Airyscan confocal microscopy. Scale bars: Main, 10 µm; ROI, 2 µm. (**B, C**) Functional assessment of filopodia formation following siRNA-mediated knockdown of specific proteins. U-87 MG cells expressing MYO10-GFP were treated with siRNAs targeting MAP4K4, MINK1, NF2, SCRIB, CSNK1A1, CCT8, or a control siRNA (siCTRL). After siRNA treatment, cells were plated on fibronectin-coated coverslips for 2 hours, then fixed and imaged using a spinning-disk confocal microscope. (**B**) Representative images for each siRNA condition. Scale bars: Main images, 20 µm; ROIs, 5 µm. (**C**) Quantification of the number of filopodia per cell. Results are shown as boxplots, with whiskers extending from the 10th to the 90th percentiles. The boxes indicate the interquartile range, and a line within each box marks the median. Data points outside the whiskers are shown as individual dots (n = 76-113 cells per condition, three biological repeats); the siCTRL condition pools data from 6 experimental sets. The numerical data and the raw images used to make this figure have been archived on Zenodo (https://zenodo.org/records/17779715).

Next, we assessed the functional roles of these proteins in filopodia formation. We performed siRNA-mediated knock-down using two independent siRNAs per candidate in U-87 MG cells expressing MYO10-GFP, and then counted the number of MYO10-positive filopodia per cell (Fig. 2B). Knockdown efficiency was confirmed by western blot analysis (Fig. S2). While silencing of MAP4K4 (85% efficiency) and NF2 (87% efficiency) did not significantly alter the number of filopodia, knocking down MINK1 (80% efficiency), CSNK1A1 (80% efficiency), SCRIB (80 to 90% efficiency), and CCT8 (75% efficiency) resulted in a significant reduction in filopodia formation (Fig. 2B-C).

This targeted screen confirms the effectiveness of our proximity-dependent biotinylation approach in identifying novel regulators of MYO10-positive filopodia. Validation of these proteins, identified in our TurboID-MYO10 datasets, as bona fide filopodia resident proteins, facilitates future investigations aimed at unraveling the complex biology of filopodia.

### CCT8 modulates filopodia independently of the TRiC complex

CCT8 (Chaperonin Containing TCP1 Subunit 8) was identified as a highly enriched protein in both U-2 OS and U-87 MG TurboID-MYO10 datasets (Fig. 1), displaying consistently high spectral counts and minimal detection in control samples (Supplementary Table 1). CCT8 is part of the TRiC (TCP1 Ring Complex), a protein-folding complex composed of eight subunits (CCT1-8) that helps fold abundant cytoplasmic proteins (Zeng et al., 2024). However, out of the eight TRiC subunits, only CCT8 was detected in our TurboID-MYO10 datasets, possibly because some TRiC sub-units have other cellular functions as monomers (Grantham, 2020). Curious about the unexpected enrichment of CCT8 in our data and its role in regulating filopodia in U-87 MG cells, we decided to investigate further.

To better understand the functional impact of CCT8 depletion, we tracked filopodia dynamics using live-cell imaging in MYO10-GFP-expressing U-87 MG cells. Filopodia in CCT8-depleted cells showed increased average speed and decreased lifetimes, indicating that CCT8 contributes to both filopodia formation and stability (Fig. S3A). Next, to assess whether CCT8’s role in filopodia is conserved, we extended our analysis beyond U-87 MG cells by targeting CCT8 expression in U-2 OS (85% efficiency) and MDA-MB-231 (85% efficiency) cells. MDA-MB-231 cells were specifically chosen because of their known reliance on MYO10-positive filopodia for migration and invasion (Arjonen et al., 2014). Consistent with findings in U-87 MG cells, knocking down CCT8 significantly reduced the number of filopodia in both U-2 OS and MDA-MB-231 cells (Fig. 3A-B and Fig. S3B-D), indicating a general role for CCT8 in regulating filopodia formation.

**Fig. 3.**
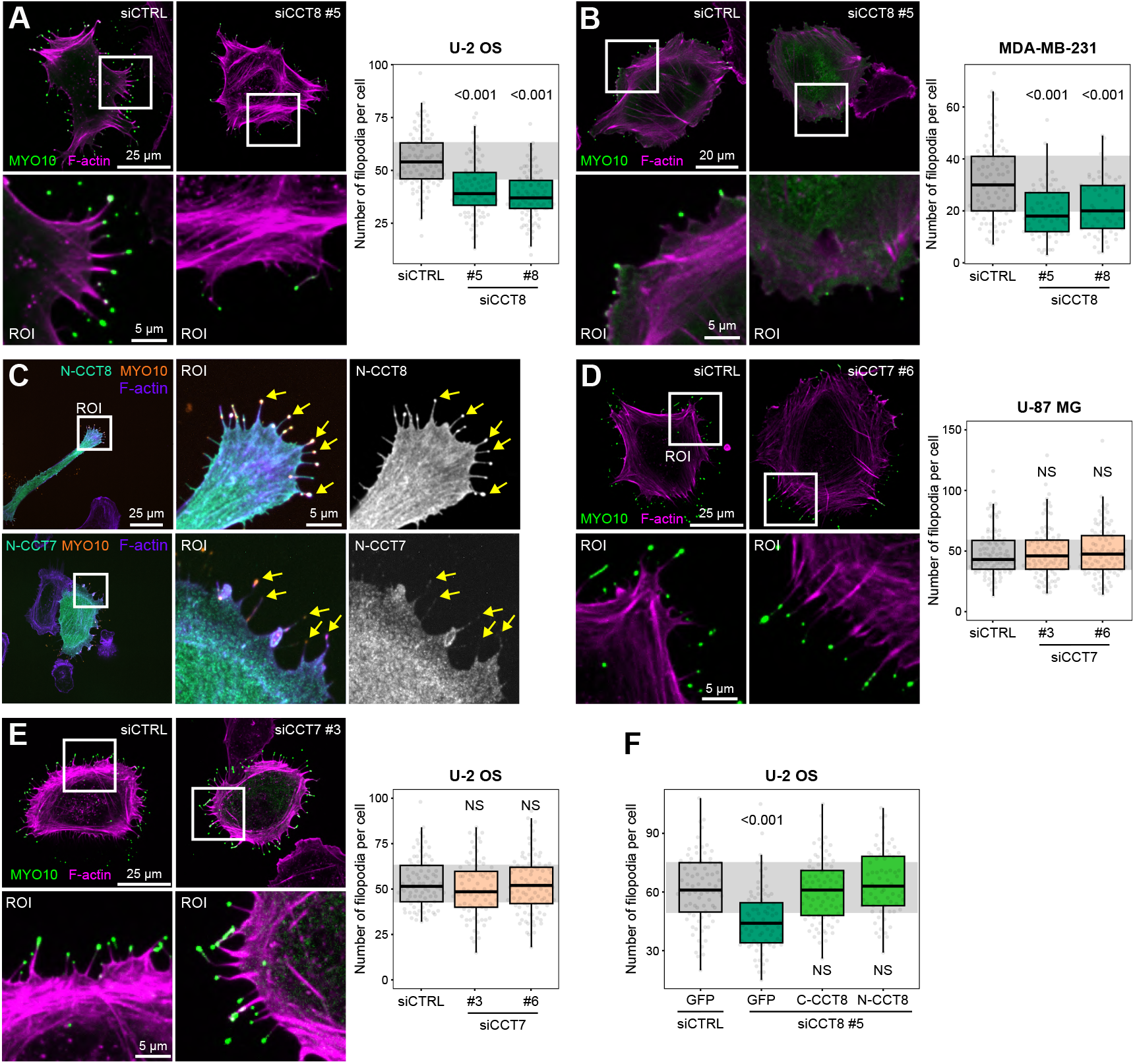
CCT8 modulates filopodia formation independently of the TRiC complex. (**A**–**B**) Filopodia quantification in U-2 OS and MDA-MB-231 cells expressing MYO10-GFP, treated with control (siCTRL) or CCT8-targeting siRNAs. Cells were plated on fibronectin-coated coverslips for 2 hours, fixed, and imaged using spinning disk confocal microscopy. (**A**) Representative images and quantification for U-2 OS cells (n > 90 cells per condition, three biological repeats). Scale bars: Main images, 25 µm; ROIs, 5 µm. (**B**) Representative images and quantification for MDA-MB-231 cells (n > 60 cells per condition, three biological repeats). Scale bars: Main images, 20 µm; ROIs, 5 µm. (**C**) Representative Airyscan confocal images of U-87 MG cells expressing MYO10-mScarlet in combination with GFP-CCT8 or GFP-CCT7. Cells were plated on fibronectin-coated coverslips for 2 hours before fixation and imaging. Scale bars: Main images, 25 µm; ROIs, 5 µm. (**D**–**E**) Filopodia quantification in U-87 MG and U-2 OS cells expressing MYO10-GFP, treated with siCTRL or CCT7-targeting siRNAs. Experimental setup as described above. (**D**) Representative images and quantification for U-87 MG cells (n > 97 cells per condition, three biological repeats). Scale bars: Main images, 25 µm; ROIs, 5 µm. (**E**) Representative images and quantification for U-2 OS cells (n > 82 cells per condition, three biological repeats). Scale bars: Main images, 25 µm; ROIs, 5 µm. (**F**) Rescue experiment examining the impact of GFP-tagged CCT8 variants on filopodia formation in CCT8-silenced U-2 OS cells. Cells expressing mScarlet-MYO10 along with GFP alone, GFP-N-CCT8 (N-terminal tagged), or GFP-C-CCT8 (C-terminal tagged) were treated with CCT8-targeting or siCTRL siRNAs, plated on fibronectin-coated coverslips for 2 hours, fixed, and imaged using spinning disk confocal microscopy. Filopodia numbers per cell were quantified. (n > 68 cells per condition, three biological repeats). (**A, B, D, E, F**) Results are shown as boxplots, with whiskers extending from the 10th to the 90th percentiles. The boxes illustrate the interquartile range, and a line inside each box indicates the median. Data points outside the whiskers are shown as individual dots. The numerical data and the raw images used to make this figure have been archived on Zenodo (https://zenodo.org/records/17779715).

To determine whether the regulation of filopodia function is specific to CCT8 or involves the TRiC complex, we first examined whether another TRiC subunit, CCT7, localizes to and/or modulates filopodia function. High-resolution imaging showed that GFP-tagged CCT7, unlike CCT8, did not accumulate at MYO10-filopodia tips (Fig. 3C). Next, we conducted similar knockdown experiments targeting CCT7 in U-87 MG (70-90% efficiency) and U-2 OS (90% efficiency) cells (Fig. 3C-D and Fig. S3E-G). Unlike CCT8, silencing CCT7 did not impact MYO10 filopodia formation (Fig. 3C-D), further supporting the idea that CCT8 regulates filopodia formation independently of the TRiC complex.

To further test this hypothesis, we used GFP-tagged CCT8 variants with tags at different ends, which, based on previous work, prevent or allow incorporation into the TRiC complex (Dekker et al., 2008; Spiess et al., 2015; Collier et al., 2021). The N-terminally tagged CCT8 (N-CCT8) is unlikely to assemble into TRiC, whereas the C-terminally tagged CCT8 (C-CCT8) should remain compatible with TRiC integration. Both C-CCT8 and N-CCT8 localized to filopodia tips (Fig. 3E), and when endogenous CCT8 was silenced, the expression of either exogenous N-CCT8 or C-CCT8 successfully rescued the number of MYO10 filopodia (Fig. 3F), further supporting a TRiC-independent mechanism. Interestingly, overexpressing either CCT8 construct in control cells did not change the number of filopodia (Fig. S3H), indicating that CCT8 is necessary for MYO10 filopodia but is not alone sufficient to induce them. Overall, these findings indicate a specific, TRiC-independent role for CCT8 in regulating MYO10-filopodia formation and dynamics.

### CCT8 binds to the MYO10 motor domain

Since CCT8 was one of the most strongly enriched proteins in our TurboID-MYO10 datasets (Fig. 1), and monomeric CCT8 regulates filopodia independently of the TRiC complex (Fig. 2 and Fig. 3), we hypothesized that CCT8 might directly interact with MYO10.

To test this, we first performed GFP-Trap pulldown assays in cells expressing either GFP-tagged full-length MYO10 or a truncated MYO10 construct, MYO10-HMM (Berg and Cheney, 2002), which includes the motor domain and the coiled-coil domains but lacks the C-terminal cargo-binding tail. Endogenous CCT8 co-precipitated efficiently with both constructs (Fig. 4A), indicating that the interaction is mediated through the N-terminal region of MYO10. This was somewhat unexpected, since the MYO10 tail, particularly the MyTH4-FERM domain, is known to mediate interactions with cargoes, including several filopodia tip proteins, such as integrins and RAPH1 (Popović et al., 2023; Wei et al., 2011).

**Fig. 4.**
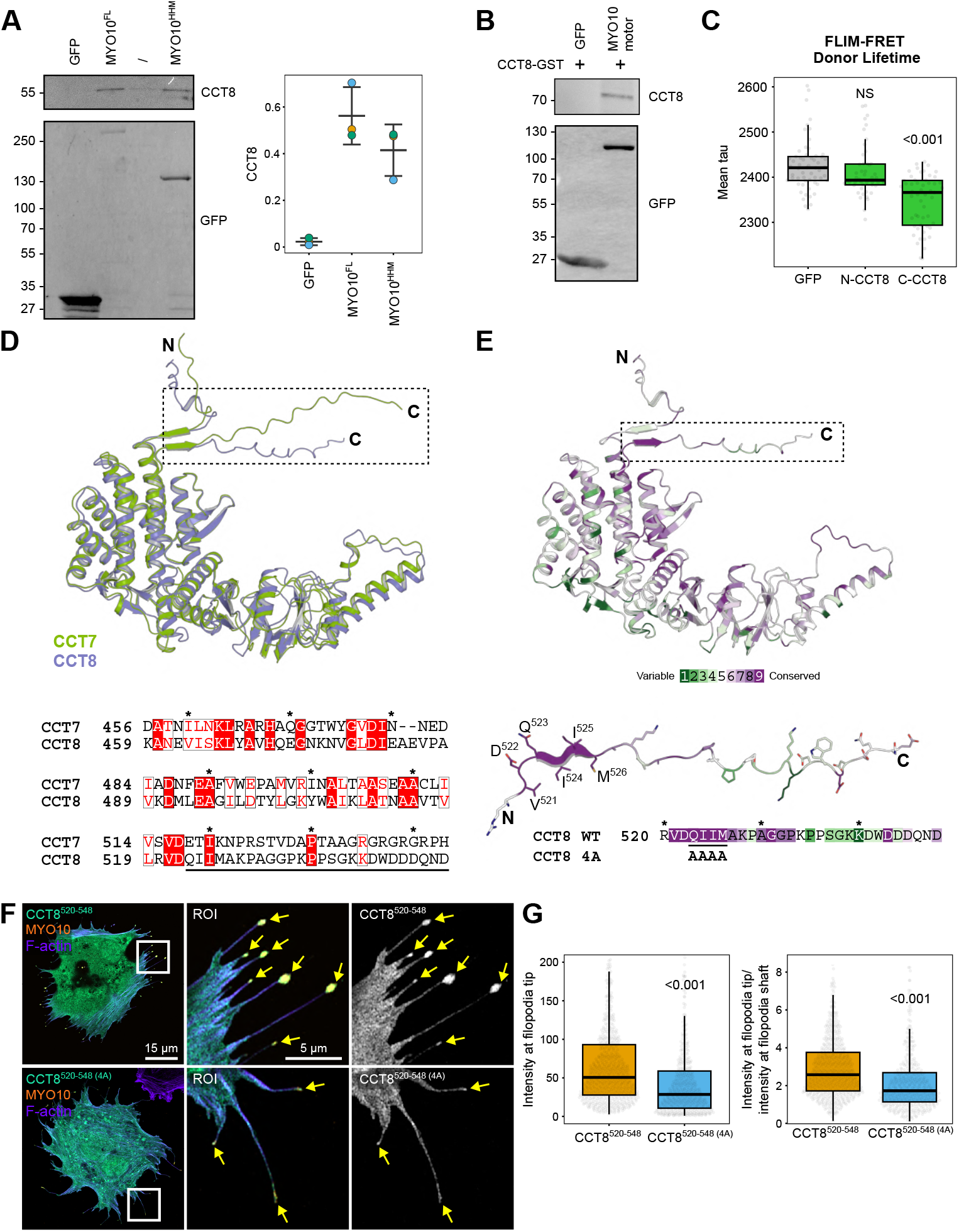
CCT8 binds to MYO10 motor domain. (**A**) GFP pulldown assays from HEK293FT cells expressing GFP alone, full-length MYO10-GFP, or the truncated MYO10 variant, MYO10-HMM. Co-precipitation of endogenous CCT8 was analyzed by western blotting. Quantifications from 3 independent experiments are presented as SuperPlots (Lord et al., 2020; Goedhart, 2021). (**B**) GFP pulldown assays indicating a direct interaction between recombinant GST-CCT8 and GFP-tagged MYO10 motor domain purified from HEK293FT cells. CCT8 binding was confirmed by western blotting. A representative western blot is displayed. (**C**) GFP lifetime (τ) was measured in U-87 MG cells co-expressing MYO10–mScarlet with GFP, GFP–CCT8 (N-terminal tag), or CCT8–GFP (C-terminal tag). The mean tau values per cell are shown (n > 49 cells per condition, four biological replicates). (**D**-**E**) The C-terminus of CCT8 features a highly conserved region. (**D**) Top: Structural alignment of CCT7 (green) and CCT8 (blue) showing a very similar tertiary structure (RMSD 1.03 Å across 2383 atoms). Bottom: sequence alignment of the last 90 residues of CCT7 and CCT8 (TCS score: 941; (Chang et al., 2014)), with the boxed region indicated by a black line. Identical residues and conserved substitutions are shown with red blocks and red text, respectively. (**E**) Top: CCT8 colored by ConSurf conservation score (Yariv et al., 2023). Bottom: detail of the boxed region illustrating residues 520-548 of CCT8 and their conservation scores. Variable residues are marked in green, neutral residues in white, and conserved residues in purple. A black line indicates the QIIM motif; the 4A mutation is indicated below. (**F**-**G**) U-87 MG cells co-expressing MYO10–mScarlet with GFP-tagged CCT8(520–548) or GFP-tagged CCT8(520–548)-4A were plated on fibronectin-coated coverslips for 2 hours before being fixed and imaged. (F) Representative Airyscan confocal images are shown. Scale bars: Main images, 15 µm; ROIs, 5 µm. (**G**) The intensity of GFP-tagged CCT8(520–548) and GFP-tagged CCT8(520–548)-4A at the filopodia tip, along with their enrichment ratio (filopodia tip/filopodia shaft), were measured using filopodia mapping (Jacquemet, 2023), see methods for details (n > 433 filopodia per condition, three biological repeats). (**C, G**) Results are shown as boxplots, with whiskers extending from the 10th to the 90th percentiles. The boxes illustrate the interquartile range, and a line inside each box indicates the median. Data points outside the whiskers are shown as individual dots. The numerical data and the raw images used to make this figure have been archived on Zenodo (https://zenodo.org/records/17779715).

To explore the CCT8-MYO10 interaction in more detail, we next tested whether the MYO10 motor domain alone can recruit CCT8. We incubated immobilized GFP-tagged MYO10 motor domain or GFP alone with recombinant GST-tagged CCT8 (CCT8-GST). CCT8-GST was pulled down only by the MYO10 motor domain, not by GFP (Fig. 4B), indicating that the MYO10 motor domain can recruit CCT8.

To verify this interaction in living cells, we used fluorescence lifetime imaging microscopy (FLIM) to measure Förster resonance energy transfer (FRET). This technique detects molecular proximity (<10 nm) between two fluorophores, providing strong evidence of a direct interaction. We co-expressed MYO10-mScarlet (FRET acceptor) with GFP, N-terminally tagged GFP-CCT8 (GFP-N-CCT8), or C-terminally tagged CCT8 (GFP-C-CCT8). A significant decrease in donor fluorescence lifetime, indicating FRET, was observed only with GFP-C-CCT8 (Fig. 4C). These findings confirm that MYO10 and CCT8 interact in living cells and further suggest that the C-terminal region of CCT8 is near MYO10’s N-terminal domain.

To identify the sequence determinants that direct CCT8 to filopodia and mediate MYO10 binding, we compared CCT8 with CCT7, which does not localize to filopodia. Although the overall protein structures are similar, their most significant differences are at the C-termini (Fig. 4D). ConSurf analysis (Yariv et al., 2023) revealed two conserved patches in CCT8: an internal loop (residues 246–274) and a C-terminal segment (residues 520–548), suggesting the C-terminus of CCT8 as a potential MYO10-interacting region (Fig. 4E).

To determine if the C-terminal region of CCT8 alone is sufficient for localizing to filopodia, we created a minimal GFP-tagged CCT8(520–548) construct. When co-expressed with MYO10, this construct accumulated at filopodia tips (Fig. 4F and Fig. 4G). Within this region, four amino acids, Gln-Ile-Ile-Met (QIIM), are highly conserved (Fig. 4D), and so we next mutated the QIIM in CCT8(524–528) to AAAA, generating the GFP-tagged CCT8(520–548)-4A mutant. Mutation of the QIIM motif reduced its accumulation at filopodia tips, further indicating that this part of CCT8 helps target filopodia and interact with MYO10 (Fig. 4F-G). Notably, in the intact TRiC complex, the 524–528 region of CCT8 is buried inside the complex (Fig. S4B), providing a structural explanation for why CCT8’s functions in TRiC and MYO10 binding are likely mutually exclusive.

### CCT8 depletion in breast cancer cells modulates cell spreading, migration, and invasion

Filopodia facilitate the spreading and migration of cancer cells by helping them sense and interact with the surrounding extracellular matrix (Jacquemet et al., 2015). Having identified CCT8 as a key regulator of MYO10-positive filopodia and given that its roles beyond TRiC complex assembly remain poorly understood, we next investigated how CCT8 depletion affects breast cancer cell behavior. MYO10 has been linked to breast cancer progression, where MYO10-positive filopodia are necessary for effective cell migration and invasion (Arjonen et al., 2014; Cao et al., 2014; Peuhu et al., 2022). Therefore, we focused our analysis on this context.

Initially, assessing the clinical significance of CCT8 in breast cancer, we examined publicly available datasets using the Kaplan-Meier plotter (Posta and Győrffy, 2025). High CCT8 expression, at both the mRNA and protein levels, was consistently associated with worse prognosis across four independent breast cancer cohorts (Fig. S5A-D). Based on this, we chose MDA-MB-231 cells, a triple-negative breast cancer cell line known for its reliance on MYO10-positive filopodia for migration and invasion (Arjonen et al., 2014), for functional studies.

We started by evaluating cell spreading on fibronectin using an impedance-based xCELLigence adhesion assay (Hamidi et al., 2017), which measures real-time changes in electrical impedance as cells attach and spread on electrode-embedded surfaces. This assay provides a dynamic readout of cell adhesion to the substrate. CCT8 depletion significantly decreased the spreading of MDA-MB-231 cells in these experiments (Fig. 5A), indicating that CCT8 contributes to effective early cell–matrix interaction.

**Fig. 5.**
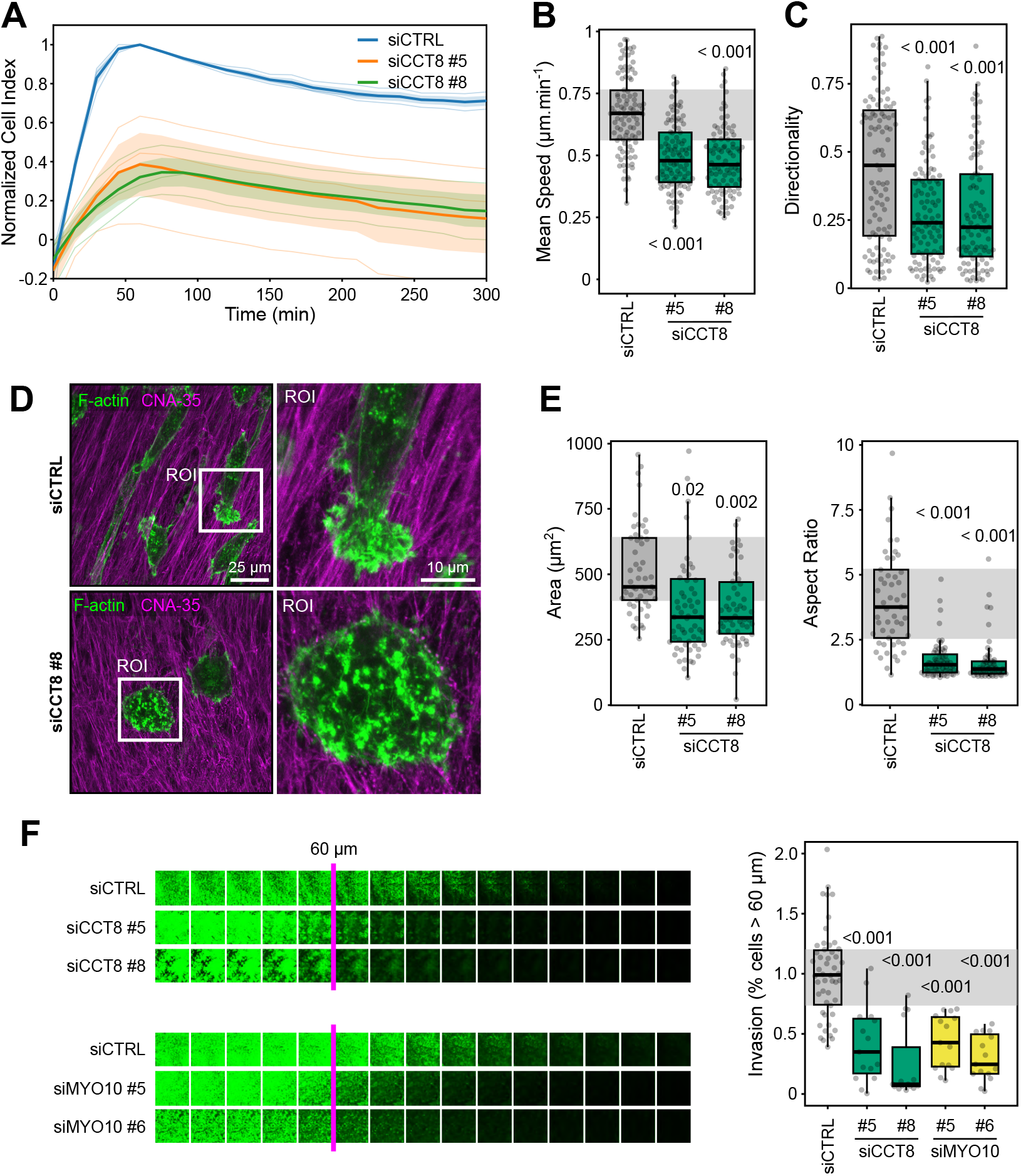
CCT8 regulates cell spreading on fibronectin, migration on fibrillar matrices, and invasion through collagen gels. (**A**) MDA-MB-231 cells were transfected with control siRNA (siCTRL) or CCT8-targeting siRNAs, then plated on fibronectin-coated wells. Spreading was monitored using xCELLigence, and data were presented as cell index over time. For each condition, thin, semi-transparent lines represent individual biological repeats (all in the same color). The bold line indicates the average for that condition, and the shaded band shows ± SEM. n = 3 biological repeats. (**B**-**E**) MDA-MB-231 cells transfected with siCTRL or CCT8-targeting siRNAs were plated on cell-derived matrices (CDMs). Cells were either live-imaged with brightfield microscopy to assess migration (**B**-**C**) or fixed, stained, and imaged with high-resolution Airyscan confocal microscopy for morphological analysis (**D**-**E**). (**B**-**C**) Quantification of migration speed for tracked cells: siCCT8 MDA-MB-231 (n > 92 tracks per condition, from three biological repeats). (**D**) Representative confocal Airyscan images of siCTRL-or siCCT8-treated MDA-MB-231 cells stained for F-actin and CNA-35 during migration on CDM. Scale bars: main images, 25 µm; insets, 10 µm. (**E**) Quantification of cell area and aspect ratio for MDA-MB-231 cells (n > 53 cells per condition, three biological repeats). (**F**) MDA-MB-231 cells transfected with siCTRL, CCT8-targeting, or MYO10-targeting siRNAs underwent an inverted transwell invasion assay using collagen I gels supplemented with fibronectin. After 48 hours, cells were fixed, stained to visualize F-actin, and imaged with confocal microscopy. Representative single optical sections, taken every 15 µm from the membrane surface, demonstrate invasion depth (shown on the left). Quantification of invasion 45 µm from the membrane is shown as boxplots on the right (n > 15 fields of view per condition, three biological repeats). (**B, C, E, F**) Results are displayed as boxplots, with whiskers extending from the 10th to the 90th percentiles. The boxes show the interquartile range, and a line inside each box indicates the median. Data points outside the whiskers are plotted as individual dots. The numerical data and the raw images used to make this figure have been archived on Zenodo (https://zenodo.org/records/17779715).

To investigate cell migration in a more physiological environment, we used cell-derived matrices (CDMs; Fig. 5B and Fig. 5C), which recapitulate the fibrillar architecture of the extracellular matrix and support filopodia-driven motility (Kaukonen et al., 2017). In MDA-MB-231 cells, CCT8 silencing resulted in a marked decrease in migration speed and directional persistence (Fig. 5B-C). High-resolution Airyscan imaging revealed that CCT8-depleted cells displayed pronounced morphological defects on CDM. In agreement with our fibronectin spreading assays, CCT8 knockdown led to smaller, more rounded cells (Fig. 5D-5E), consistent with impaired protrusive activity.

We next examined the impact of CCT8 depletion on invasion using an inverted 3D collagen invasion assay. In this setup, cells are seeded on the underside of collagen-coated transwell inserts and allowed to invade through the collagen matrix toward a serum-containing chemoattractant. As expected, MYO10 depletion (92% knockdown efficiency) significantly reduced the ability of MDA-MB-231 cells to invade through 3D collagen matrices (Fig. 5F). Notably, CCT8 depletion similarly decreased MDA-MB-231 invasion (Fig. 5F), mimicking the effects of MYO10 loss.

Together, these data demonstrate that CCT8 is crucial for effective breast cancer cell spreading, migration on fibrillar matrices, and invasion. The substantial phenotypic similarity between CCT8 and MYO10 depletion supports a model in which CCT8 promotes cell adhesion and migration, at least partly, by regulating MYO10-positive filopodia. Additionally, CCT8 inhibition is likely to cause broader, filopodia-independent effects, potentially through its TRiC-related functions, which further enhance these defects.

## Discussion

Here, we used TurboID-mediated proximity labeling to compile a comprehensive list of hundreds of proteins associated with MYO10-positive filopodia tips and with MYO10 itself. From these TurboID-MYO10 datasets, we identified MINK1, SCRIB, CSNK1A1, and CCT8 as new regulators of MYO10 filopodia. Focusing on CCT8, we show that it regulates MYO10 function independently of the TRiC complex and interacts with the MYO10 motor domain. Functionally, CCT8 influences cell spreading, migration, and invasion in breast cancer cells.

Using TurboID allowed us to greatly expand the variety and detail of MYO10-proximal proteins identified compared to our previous MYO10-BioID dataset (Popović et al., 2023), likely because TurboID has faster reaction kinetics. Our TurboID-MYO10 dataset revealed many known filopodia regulators, including FMNL2 (Pfisterer et al., 2020), RAPH1/lamellipodin (Krause et al., 2004), ITGB1 (Zhang et al., 2004), TLN1 (Jacquemet et al., 2016; Lagarrigue et al., 2015), and LASP1 (Jacquemet et al., 2019). In addition to these well-known filopodial components, the proximity proteomes highlight a broad network of cortical and junctional scaffolds (TLN2, ZYX, EPB41L2, AHNAK2, TJP1/2, SCRIB, NF2, and TAGLN/TAGLN2), signaling molecules, Rho GTPase regulators (e.g., CDC42BPA, PREX1, ARHGEF7, ARHGAP10/22), and vesicle trafficking factors (SYTL4, RAB11FIP2/5, AP3S1/2, RAB27B, ERBB2IP, STON2). Importantly, we also identified multiple regulators of membrane lipids and phosphoinositides, including PIP4K2A/C, PIP5K1C, MTMR2, SYNJ1/2, oxysterol-binding proteins (OSBPL6/7/8), and enzymes involved in fatty acid and phospholipid metabolism (FASN, LYPLA2, CEPT1, PGAP1), along with lipid-binding scaffolds (ANXA1/2, FABP5, PHLDB1). Overall, these findings support the idea that MYO10 functions within phosphoinositide- and lipid-rich cortical microdomains at the plasma membrane, and that their regulation is vital for filopodia activity (Jacquemet et al., 2019; Mason et al., 2025; Plantard et al., 2010). However, some known filopodial components, including p130Cas (Jacquemet et al., 2019) and VASP (Lanier et al., 1999), were not detected, perhaps due to the spatial limitations of proximity labeling. Interestingly, the overlap between the U-87 MG and U-2 OS TurboID-MYO10 datasets was limited, indicating that MYO10’s proximal network may vary significantly between cell types. Overall, our TurboID-MYO10 approach and datasets will be valuable tools and resources for researchers studying the composition and organization of MYO10 filopodia.

We identified CCT8 as a MYO10-associated factor, indicating functions for CCT8 outside the traditional TRiC chaperonin cycle. This finding supports the growing evidence that individual CCT subunits can operate as monomers or as part of subcomplexes, performing roles beyond protein folding (Grantham, 2020). Such “moonlighting” activities have been observed for several CCT subunits. For example, CCT2 acts as an autophagy receptor, recruiting ubiquitinated cargo for degradation (Wang et al., 2025; Ma et al., 2022). CCT3 can protect cells from drug-induced ferroptosis by hindering the recycling of transferrin receptor protein 1 (Zhu et al., 2024). Similarly, overexpression of CCT4 promotes filopodia formation independently of TRiC-mediated folding (Spiess et al., 2015). Our findings extend this concept to CCT8, although the precise molecular mechanism by which CCT8 influences MYO10 function remains unknown. Based on our mapping and modeling, possible scenarios include: (i) changing MYO10’s motor activity or duty ratio, thereby affecting its processivity, or (ii) influencing MYO10’s activation and dimerization. Future work will aim to decipher the exact mechanism by which CCT8 modulates MYO10 function. It is also important to note that we cannot exclude the possibility that some of the phenotypes we observed are further amplified by CCT8’s role in modulating the cytoskeleton (Oftedal et al., 2021; Kelly et al., 2022; Yang et al., 2018).

Filopodia have long been recognised as regulators of cancer cell migration and invasion across various cancer types (Jacquemet et al., 2015). Here, we identify CCT8 as a modulator of MYO10-positive filopodia in cancer cell lines from different origins, including osteosarcoma, glioma, and breast cancer. Since the roles of filopodia are particularly well understood in breast cancer, we focused on this context. We found that CCT8 enhances breast cancer cell adhesion, migration, and invasion through collagen gels. Consistent with these functional findings, high CCT8 expression is linked with poor outcomes in breast cancer, aligning with previous pan-cancer analyses (Gong et al., 2023). Additionally, earlier studies have shown that CCT8 knockdown hampers migration in glioma cells by disrupting actin organization (Qiu et al., 2015) and that CCT8 promotes invasion in esophageal squamous cell carcinoma and colorectal cancer by affecting cytoskeletal and EMT-related pathways (Yang et al., 2018; Liao et al., 2021). Collectively, these findings support a broader role for CCT8 in promoting cancer migration and invasion. Given its influence on breast cancer cell behavior and promising clinical data, future research will explore the role of CCT8 in breast cancer progression using preclinical approaches, including in vivo models.

## Methods

### Cells

HEK293 (human embryonic kidney cells), MDA-MB-231 (triple-negative human breast adenocarcinoma), TIFFs (human telomerase-immortalized foreskin fibroblasts), and U-2 OS (human bone osteosarcoma epithelial cells) were cultured in DMEM (Dulbecco’s Modified Eagle’s Medium; Sigma, D1152) supplemented with 10% fetal bovine serum (FBS) (Biowest, S1860). U-87 MG glioblastoma cells were cultured in MEM (Minimum Essential Medium; Gibco, A4192201) with 10% FBS (Biowest, S1860). U-2 OS cells were obtained from DSMZ (Deutsche Sammlung von Mikroorganismen und Zellkulturen, Braunschweig, Germany, ACC 785). MDA-MB-231 and U-87 MG cells were provided by ATCC (HTB-26 and HTB-14). HEK293 cells were supplied by ATCC (CRL-1573). TIFFs were donated by Jim Norman (CRUK, Beatson Institute, UK). All cells were tested for mycoplasma contamination. Cell lines have not been authenticated.

Stable U-87 MG and U-2 OS cell lines expressing MYO10-GFP or MYO10–TurboID were created through transient transfection with the appropriate expression plasmids. Twenty-four hours after transfection, cells were selected using Geneticin (G418; Gibco, 11811098) at a final concentration of 1 mg/mL. Successfully transfected cells were maintained in complete medium supplemented with 1mg/mL G418 to sustain selection pressure during routine culture. MYO10-TurboID-expressing cells were cultured in a medium containing 100 µM avidin to reduce background biotinylation.

### Antibodies and reagents

The following primary antibodies were used in this study for immunofluorescence (IF) and western blotting (WB). Rabbit antibodies: anti-MYO10 (PA5-55019, Thermo Fisher Scientific; WB 1:1000), anti-MAP4K4 (ab80418, Abcam; WB 1:1000), anti-MINK1 (PA5-288901, Sigma-Aldrich; WB 1:1000), anti-SCRIB (HPA064312, Sigma-Aldrich; WB 1:1000), anti-CCT7 (PA5-98811, Invitrogen; WB 1:1000), and anti-NF2 (HPA003097, Sigma-Aldrich; WB 1:1000). Mouse antibodies: anti-GAPDH (5G4 / 5G4cc, Hytest; WB 1:2000), anti-CCT8 (sc-377453, Santa Cruz Biotechnology; WB 1:1000), anti-CSNK1A1 (sc-74582, Santa Cruz Biotechnology; WB 1:1000), and anti-α-tubulin (12G10, DSHB; WB 1:2000).

For F-actin staining, phalloidin conjugated to Alexa Fluor 488 (A12379, Thermo Fisher Scientific), Alexa Fluor 555 (A34055, Thermo Fisher Scientific), or Alexa Fluor 647 (A30107, Thermo Fisher Scientific) was used at 1:2000 for IF. Biotinylated proteins were detected using streptavidin-Alexa Fluor 555 (S11223, Thermo Fisher Scientific; IF 1:200) or streptavidin-Alexa Fluor 680 (S21378, Thermo Fisher Scientific; IF 1:200, WB 1:500). Collagen was visualized using CNA-35-GFP (Aper et al., 2014), produced in-house.

### Plasmids

MYO10-mScarlet was previously generated (Jacquemet et al., 2019) and is available from Addgene (plasmid #145179). The MYO10-GFP (EGFPC1-hMyoX) plasmid was a gift from Emanuel Strehler (Addgene plasmid 47608) (Bennett et al., 2007). The MYO10HHM-GFP plasmid (boundaries 2-948 aa from full-length bovine MYO10) was a kind gift from Timothy Andrew Sanders (University of Chicago, USA). The CCT8-Nter-GFP (CCT8-pcDNA3.1(+)-N-eGFP), CCT8-Cter-GFP (CCT8-pcDNA3.1(+)-C-eGFP), and MYO10motor-GFP (boundaries 1-741 aa from full-length human MYO10) MYO10_MD_pcDNA3.1(+)-N-eGFP were purchased from Genscript. Briefly, the gene fragments were synthesized using gene synthesis and cloned into pcDNA3.1(+)-N-eGFP at the BamHI/XhoI (CCT8-Nter-GFP, MYO10motor-GFP) or into pcDNA3.1(+)-C-eGFP at the NheI/NotI (CCT8-Cter-GFP) sites. All constructs were sequence-verified. The TurboID-MYO10 plasmid was generated by replacing the EGFP sequence in the MYO10-GFP plasmid with the TurboID sequence (Branon et al., 2018) using a gene block. The CCT8-GST construct was generated by cloning the CCT8 sequence from the CCT8-pcDNA3.1(+)-N-eGFP plasmid into the pGEX 4T1 vector. The GFP-CCT8 insert was obtained by KpnI digestion, blunted, then cut with XhoI and ligated into the SmaI/XhoI-digested pGEX 4T1 backbone. The CCT8(520–548) WT and 4A mutant constructs were synthesised by GeneArt (Life Technologies) and sub-cloned into pEGFP-C1 (with a modified multiple-cloning site based on pET47b) between the XmaI/SacI sites using ligation-independent cloning.

Several of the GFP-tagged constructs used here were generated by the Genome Biology Unit core facility cloning service (Research Programs Unit, HiLIFE Helsinki Institute of Life Science, Faculty of Medicine, University of Helsinki, Biocenter Finland) by transferring entry clones from the ORFeome collaboration library into mEmerald destination vectors using the standard LR reaction protocol. The entry clones (I.M.A.G.E. Consortium CloneID) transferred into pcDNA6.2/N-EmGFP-DEST include MAP4K4 (100062694), CCT7 (100006094), and NF2 (100000483). The entry clones (I.M.A.G.E. Consortium CloneID) transferred into pcDNA6.2/C-EmGFP-DEST include MINK1 (100003788), SCRIB (100015198), and CCT7 (100010794). pDONR223_CSNK1A1_WT was a gift from Jesse Boehm, William Hahn, and David Root (Addgene plasmid 82106) (Kim et al., 2016) and was transferred into pcDNA6.2/N-EmGFP-DEST using the standard LR reaction protocol.

### Plasmid transfection and siRNA-mediated gene silencing

U-2 OS and U-87 MG cells were transfected with the indicated plasmids using Lipofectamine 3000 and the P3000 Enhancer Reagent (Thermo Fisher Scientific, L3000001), following the manufacturer’s instructions. Protein expression was suppressed using 100 nM siRNA and Lipofectamine 3000 (Thermo Fisher Scientific) following the manufacturer’s instructions. The siRNAs used for silencing were:

- **CCT8**: siCCT8#5 (Hs_CCT8_5; SI03026212), siCCT8#8 (Hs_CCT8_8; SI03117261), siCCT8#6 (Hs_CCT8_6, SI03036460), siCCT8#7(Hs_CCT8_7, SI03110681).
- **MYO10**: siMYO10#5 (Hs_MYO10_5; SI04158245), siMYO10#6 (Hs_MYO10_6; SI04252822).
- **MAP4K4**: SI02223704), siMAP4K4#5 (Hs_MAP4K4_5, siMAP4K4#6 (Hs_MAP4K4_6, SI02223711) siMAP4K4#9 (Hs_MAP4K4_9, SI02633911), siMAP4K4#11 (Hs_MAP4K4_11, SI03115098).
- **MINK1**: siMINK1#3 (Hs_MINK1_3, SI00112133), siMINK1#10 (Hs_MINK1_10, SI04954978), siMINK1#4 (Hs_MINK1_4, SI00112140) siMINK1#6(Hs_MINK1_6, SI02759022).
- **SCRIB**: siSCRIB#6 (Hs_SCRIB_6, SI04182290), siSCRIB#7 (Hs_SCRIB_7, SI04295655), siSCRIB5(Hs_SCRIB_5, SI03157700), CRIB1(Hs_SCRIB_1, SI00712635). siS-
- **NF2**: siNF2#7 (Hs_NF2_7, SI02664137), siNF2#17 (Hs_NF2_17, SI04949952), siNF2#8 (Hs_NF2_8, SI02664452), siNF2#18 (Hs_NF2_18, SI04949959).
- **CSNK1A1**: siCSNK1A1#5 (Hs_CSNK1A1_5, SI00605395), siCSNK1A1#11 (Hs_CSNK1A1_11, SI04434829); siCSNK1A1#6 (Hs_CSNK1A1_6, SI00605402), siCSNK1A1#10 (Hs_CSNK1A1_10, SI04434822).
- **CCT7**: siCCT7#3 (Hs_CCT7_3, SI00340207), siCCT7#6 (Hs_CCT7_6, SI03236737)

Control cells were silenced using AllStars Negative Control siRNA (SI03650318), with all siRNAs provided by Qiagen.

### SDS–PAGE and quantitative western blotting

Protein extracts were separated by SDS-PAGE under denaturing conditions and transferred to nitrocellulose membranes using a Mini Blot Module (Invitrogen, B1000). Membranes were blocked for 30 minutes at room temperature with 1× StartingBlock buffer (Thermo Fisher Scientific, 37578). After blocking, the membranes were incubated overnight with the appropriate primary antibody (1:1000 in blocking buffer), washed three times with PBS, and then probed for 1 hour with a fluorophore-conjugated secondary antibody diluted 1:5000 in blocking buffer. The membranes were washed three times with PBS for 30 minutes and scanned using an iBright FL1500 imaging system (Invitrogen).

### GFP-trap pull-down

Cells transiently expressing the indicated GFP-tagged bait proteins were lysed in ice-cold lysis buffer containing 25 mM Tris-HCl (pH7.4), 150 mM NaCl, 1 mM EDTA, 1%(v/v) NP-40, and protease inhibitor cocktail (Pierce Protease Inhibitor Mini Tablets, A32953, ThermoFisher). Lysates were clarified by centrifugation at 15,000 g for 10 minutes at 4°C. The supernatant was incubated with GFP-Trap magnetic beads (gtma-20, ChromoTek) for 2 hours at 4°C with gentle rotation. Beads were collected by magnetic separation and washed three times with a lysis buffer. The bound proteins were then eluted in Laemmli sample buffer by boiling for 10 minutes.

For in vitro binding assays testing the interaction between the MYO10 motor domain and recombinant CCT8, lysates containing GFP-tagged MYO10 motor domain were incubated with GFP-Trap beads as described above. This was followed by three washes with a high-stringency buffer (1% (v/v) Triton X-100, 1 M NaCl, 100 mM Tris-HCl, pH 6.5, 10 mM DTT). The beads were then incubated overnight at 4°C with purified GST-CCT8. After three washes with lysis buffer, protein complexes were eluted in Laemmli sample buffer and analyzed by SDS-PAGE and western blotting.

### Proximity biotinylation

U-2 OS and U-87 MG cells stably expressing MYO10–TurboID, previously grown in biotin-free medium, were plated on fibronectin-coated plates. After 1 hour, the biotin-free medium was replaced with a medium containing either 50 µM biotin or 100 µM avidin, and the cells were incubated for an additional hour. After washing with cold PBS, the cells were lysed, and debris were removed by centrifugation (15,000 g, 4°C, 10 min). Biotinylated proteins were then incubated with streptavidin beads (MyOne Streptavidin C1, Invitrogen) for 1 hour with rotation at 4°C. Beads were washed twice with 500µL wash buffer 1 (10% (w/v) SDS), once with 500µL wash buffer 2 (0.1% (w/v) deoxycholic acid, 1% (v/v) Triton X-100, 1mM EDTA, 500mM NaCl, and 50mM HEPES), and once with 500µL wash buffer 3 (0.5% (w/v) deoxycholic acid, 0.5% (w/v) NP-40, 1mM EDTA, and 10mM Tris/HCl pH 7.4). Proteins were eluted in 70µL of 2× reducing sample buffer (100 mM Tris-HCl, pH 6.5, 4% (w/v) SDS, 17.5% (v/v) glycerol, 3 mM bromophenol blue, 0.2 M dithiothreitol) for 10min at 95°C.

### Mass spectrometry analysis

Affinity-captured proteins were separated by SDS-PAGE and allowed to migrate 5 mm into a 4–12% polyacrylamide gel. After staining with InstantBlue (Expedeon), gel lanes were sliced into two 2-mm bands. The slices were washed three times with water, then washed twice with a solution of 0.04 M ammonium bicarbonate and 50% acetonitrile until all the blue dye disappeared. Gel slices were shrunk with 100% acetonitrile for 5–10 minutes and rehydrated in a reducing buffer containing 20 mM dithiothreitol in 100 mM ammonium bicarbonate for 30 minutes at 56°C. Proteins within the gel pieces were then alkylated by washing the slices with 100% acetonitrile for 5–10 minutes, followed by rehydration in an alkylating buffer of 55 mM iodoacetamide in 100 mM ammonium bicarbonate solution (protected from light) for 20 minutes at room temperature. Finally, the gel pieces were washed twice with 100 mM ammonium bicarbonate. The slices were dehydrated with 100% acetonitrile and thoroughly dried using a vacuum centrifuge. Gel pieces were incubated in trypsin solution (0.02 µg/µL; Promega, cat. no. V5111) on ice for 20 minutes. After this, a solution of 40 mM ammonium bicarbonate with 10% acetonitrile was added, and protein digestion was continued overnight at 37°C. Following trypsin digestion, an equal volume of 100% acetonitrile was added, and the gel pieces were incubated again at 37°C for 15 minutes. Peptides were then extracted using a buffer of 50% acetonitrile and 5% formic acid. The peptide-containing buffer was collected, and the sample was dried in a vacuum centrifuge. The dried peptides were stored at 20°C. Before LC-ESI-MS/MS analysis, dried peptides were dissolved in 0.1% formic acid.

The LC-ESI-MS/MS analyses were performed using a nanoflow HPLC system (Easy-nLC 1200, Thermo Fisher Scientific) coupled to a Q Exactive HF mass spectrometer (Thermo Fisher Scientific, Bremen, Germany) equipped with a nano-electrospray ionization source. Peptides were first loaded onto a trapping column and then separated online on a 15 cm C18 column (75 µm x 15 cm, ReproSil-Pur 3 µm 120 Å C18-AQ, Dr. Maisch HPLC GmbH, Ammerbuch-Entringen, Germany). The mobile phase consisted of water with 0.1% formic acid (solvent A) or acetonitrile/water (80:20 (v/v)) with 0.1% formic acid (solvent B). A linear gradient over 20 minutes, from 6% to 39% B, was used to elute the peptides. MS data were acquired automatically using Thermo Xcalibur 4.1 software (Thermo Fisher Scientific). The information-dependent acquisition method involved an Orbitrap MS survey scan of the mass range 350–1750 m/z, followed by HCD fragmentation of the ten most intense peptide ions.

Raw data from the mass spectrometer were processed using Proteome Discoverer 2.4 (Thermo Fisher Scientific) and submitted to the Mascot search engine. The search was conducted against the human SwissProt database (version 2021_02), assuming trypsin as the digestion enzyme and a maximum of 2 missed cleavages. The initial mass tolerance was set to 10 ppm (parts per million) for precursor ions, and the fragment ion mass tolerance was 0.02 Dalton. Cysteine carbamidomethylation was set as a fixed modification, while methionine oxidation and N-terminal acetylation were treated as variable modifications.

To generate the U-87 MG TurboID-MYO10 dataset, four biological replicates were combined. To create the U-2 OS TurboID-MYO10 dataset, three biological replicates were combined. Proteins enriched at least 1.5-fold in biotin-supplemented TurboID-MYO10 compared to control TurboID-MYO10 (supplemented with avidin) (based on spectral counts) and detected with over six spectral counts (across all repeats) were considered potential MYO10-binding proteins. The fold-change enrichment and the significance (Saint Score) of the association were calculated using CRAPome 2.0 (Mellacheruvu et al., 2013) to generate the volcano plots (Fig. 1). Fold change was calculated by geometrically averaging replicates with three virtual controls. The Saint Score was obtained by averaging the replicate results using 10 virtual controls (n-burn 2000, n-iter 4000, LowMode-0, MinFold-1, Normalize-1).

Gene Ontology analysis was conducted using ShinyGO (v 0.82) (Ge et al., 2020). P-values were calculated with the hypergeometric test, and false discovery rates (FDRs) were determined via the Benjamini-Hochberg method to correct for multiple comparisons. Fold enrichment is defined as the percentage of genes in the list that are part of a pathway, divided by the corresponding percentage in the background. Expression levels were obtained from the Cancer Cell Line Encyclopedia (CCLE) (version DepMap Public 22Q2). Expression data are shown as RNA-seq TPM gene expression, calculated using RSEM and log2-transformed, with a pseudo-count of 1.

### Light microscopy setup

The spinning-disk confocal microscope used was a Marianas spinning-disk imaging system with a Yokogawa CSU-W1 scanning unit on an inverted Zeiss Axio Observer Z1 microscope controlled by SlideBook 6 (Intelligent Imaging Innovations, Inc.). Images were acquired using either an Orca Flash 4 sCMOS camera (chip size 2,048 × 2,048; Hamamatsu Photonics) or an Evolve 512 EMCCD camera (chip size 512 × 512; Photometrics). The objectives used were 63×/1.4 NA O Plan-Apochromat (Zeiss) and 100×/1.4 NA O Plan-Apochromat (Zeiss)

The structured illumination microscope (SIM) used was DeltaVision OMX v4 (GE Healthcare Life Sciences), fitted with a 60× Plan-Apochromat objective lens, 1.42 NA (immersion oil RI of 1.516), operating in SIM illumination mode (five phases × three rotations). Emitted light was collected on a front-illuminated pco.edge sCMOS (pixel size 6.5 nm, readout speed 95 MHz; PCO AG) controlled by SoftWorx. The confocal microscope used was a laser scanning confocal microscope (LSM880, Zeiss), equipped with an Airyscan detector (Carl Zeiss) and a 40× water (NA 1.2) or 63× oil (NA 1.4) objective. The microscope was controlled using Zen Black (2.3), and the Airyscan was used in standard super-resolution mode.

The FLIM confocal microscope used was a Leica Stellaris 8 Falcon, equipped with an 86× water-immersion objective (NA 1.2). The microscope was controlled by LAS X software (4.8.2).

### FLIM-FRET Analysis

U-87 MG cells transiently expressing MYO10-mScarlet (acceptor) and EGFP, C-terminus EGFP-tagged CCT8, or N-terminus EGFP-tagged CCT8 (donors) were plated on fibronectin-coated Ibidi 8-well glass-bottom dishes for two hours at 37°C. Cells were fixed with 4% PFA and imaged on a Leica Stellaris 8 Falcon FLIM confocal microscope using an 86x water-immersion objective. A white-light laser provided excitation at 40 MHz and 488 nm. Fluorescence was recorded using a hybrid detector in photon-counting mode, with emission collected over 495–530 nm.

Fitting of FLIM images was performed with the FLIMfit software tool (version 5.0.3). Images were spatially binned 2×2 to ensure sufficient photons per pixel before data fitting. The fitting was performed over a time window ranging from 200 ps to 15,000 ps. A minimum intensity threshold of 100 counts was applied to exclude low-signal pixels from the analysis, while the maximum gate was limited to 1 × 10^8^ counts. The instrument response function was measured using a nanogold solution (Sigma-Aldrich, 752584).

### Filopodia formation assays on fibronectin

Cells expressing human MYO10-GFP were plated for 2 hours on glass-bottom 8-well dishes coated with 10µg/mL of fibronectin. The cells were then fixed using 4% PFA, washed with PBS, and stained with phalloidin. To determine the number of filopodia, images were captured with a spinning-disk confocal microscope using a 100× objective, and filopodia per cell were manually counted.

### Filopodia dynamic assay

To study filopodia stability, U-87 MG cells expressing MYO10–GFP were plated on fibronectin-coated 35 µm glass-bottom dishes (ibidi, 81218) for at least two hours before live imaging began. Images were captured every 5 seconds at 37°C using an Airyscan microscope with a 63× objective. All live-cell imaging experiments were conducted in standard growth medium supplemented with 50 mM HEPES at 37°C under 5% CO_2_. Filopodia lifetimes were measured by identifying and tracking all MYO10 spots with the Fiji plugin TrackMate (Ershov et al., 2022). In TrackMate, the custom Stardist detector and the LAP tracker (Linking max distance = 1 micron, Gap-closing max distance = 0 microns, Gap-closing max frame gap = 0 microns) were used. The StarDist 2D model was trained for 200 epochs on 11 paired image patches (image dimensions: (512, 512), patch size: (512,512)) with a batch size of 2, using a mean absolute error (MAE) loss function. Training utilized the StarDist 2D Zero-CostDL4Mic notebook (von Chamier et al., 2021; Schmidt et al., 2018) and was accelerated with a Tesla K80 GPU.

### Mapping of CCT8 constructs within filopodia

To map the localization of each CCT8 construct within filopodia, images were first processed in Fiji and data analyzed using R as previously described (Jacquemet, 2023). Briefly, in Fiji (Schindelin et al., 2012), the brightness and contrast of each image were automatically adjusted, with the brightest cellular structure within the field of view serving as the upper limit. In Fiji, line intensity profiles (1-pixel width) were manually drawn from the filopodium tip to the base (defined as the point of intersection between the filopodium and the lamellipodium). To prevent bias in the analysis, the intensity profile lines were drawn from a merged image of actin, MYO10, and CCT8. All visible filopodia in each image were analyzed and exported for further analysis (using the “Multi Plot” function). Line intensity profiles were then compiled and analyzed in R. To standardize filopodia length, each line intensity profile was divided into 40 bins (using the median pixel value in each bin and the R function “tapply”). The filopodium tip was defined as bins 1-6, and the filopodia shaft as bins 7-40, based on MYO10 profiles.

### Cell migration on cell-derived matrices

Cell-derived matrices (CDMs) were prepared as previously described (Kaukonen et al., 2017). Briefly, TIFFs were seeded at a density of 50,000 cells per mL in a 24-well plate. Once confluent, cells were cultured for an additional 7 days, with the medium changed every 48 hours and supplemented with complete medium containing 50 mg/mL ascorbic acid (Sigma-Aldrich, A92902) to promote collagen cross-linking. Mature matrices were then stripped of cells using a lysis buffer (PBS containing 20 mM NH4OH and 0.5% (v/v) Triton X-100). After PBS washes, matrices were incubated with 10 mg/mL DNase I (Roche, 10104159001) at 37°C for 30 minutes. Finally, matrices were stored in PBS with 1% (v/v) penicillin/streptomycin at 4°C until use. For cell migration analyses, U-87 MG or MDA-MB-231 cells were seeded on CDMs for 4 hours before imaging. Brightfield images were taken every 10 minutes over 16 hours using a CellIQ system with a 10× objective at 37°C and 5% CO_2_. Cell tracking was performed manually using Fiji’s Manual Tracking plugin to assess cell migration (Schindelin et al., 2012). Only cells that met the following criteria were included: (i) not dividing during the imaging period, (ii) not colliding with other cells, and (iii) not completely immobile during the imaging period. Migration parameters were calculated from the tracking data and compiled using CellTracksColab (Gómez-de Mariscal et al., 2024). For morphology quantification, cells were seeded on CDMs overnight and fixed with 4% PFA. Cells were visualized with phalloidin, and CNA35-GFP was used to visualize the matrix (Aper et al., 2014). Samples were imaged on a confocal microscope (LSM880; Zeiss). Cells were manually segmented, and area, aspect ratio, and circularity were measured in Fiji (Schindelin et al., 2012).

### Adhesion assay on RTCA system

Plates containing gold microelectrodes (E-Plate VIEW 96, 300601020, Agilent) were coated with 10 µg/mL fibronectin in PBS for 1 hour at 37°C. Plates were washed with PBS, and 15000 U-87 MG or MDA-MB-231 cells were seeded into each well. Cellular impedance was recorded with xCELLigence RTCA eSight every 15 minutes for 6 hours at 37°C and 5% CO_2_. Cell index was determined using the xCELLigence RTCA software system. All replicate traces were normalised to the maximum cell index of the siCTRL condition within the same experiment (across time), setting siCTRL_max = 1 and expressing all values as a fraction of this reference. Plots were generated in Python using Pandas, NumPy, and Matplotlib.

### Invasion assay

Transwell inserts (8µm ThinCert; Greiner) were filled with 200µL of collagen I (concentration 5µg/mL; PureCol EZ Gel, Advanced BioMatrix) supplemented with 10 µg/mL fibronectin and allowed to polymerize for 1hour at 37°C. The inserts were then inverted, and cells were seeded on the bottom of each insert. Transwell inserts were placed in a well containing serum-free medium, with medium supplemented with 10% FBS added on top of the collagen gel. Cells were fixed with 4% PFA 48 hours after seeding and stained with phalloidin-Alexa Fluor 488. Inserts were washed three times with PBS and imaged on a confocal microscope (LSM880; Zeiss). Serial optical sections were captured every 15µm for 210 µm using a 10× objective lens. Invasion was quantified in Fiji (Schindelin et al., 2012) by measuring, for each field of view, the area occupied by cells located 60µm from the membrane surface, expressed as a percentage of the total cell-covered area within the image stack.

### Production of GST-CCT8

The GST-CCT8 protein was produced by transforming the recombinant plasmid GST-CCT8 into E. coli strain BL21(DE3)Star. Cells were cultured in 2× TY medium in ten 500 mL flasks at 37°C until the optical density at 600 nm reached 0.6. The cells were then induced with 1 mM IPTG and incubated overnight at 21°C with shaking at 175 rpm. Cells were harvested by centrifugation, yielding 37.7g of wet cell mass, and resuspended in lysis buffer (25mM Hepes, 250mM NaCl, 1mg/mL lysozyme, 1× protease inhibitor cocktail, pH 7.0). Cell disruption was performed using an Avestin Emulsiflex C3 homogenizer, and cellular debris were removed by centrifugation at 75,000 × g for 45 min. The cleared lysate was incubated with 5 mL of Pierce GSH-Agarose resin at 4°C for 30 minutes with agitation. The resin was washed with buffer (25mM Hepes, 250mM NaCl, pH 7.0), and bound proteins were eluted using GSH elution buffer (25mM Hepes, 250mM NaCl, 10mM GSSG, pH 7.0). Eluted fractions were analyzed by SDS-PAGE, and protein content was further assessed by size exclusion chromatography (Superdex 200 Increase 10/300 GL).

### Statistical analysis

The code for performing randomization tests and t-tests is available on GitHub (https://github.com/CellMigrationLab/Plot-Stats). Randomization tests used Cohen’s d as the effect size metric, with 10,000 iterations for each test. For t-tests, data were assumed to follow a normal distribution, although this was not formally tested.

## Data availability

Plasmids generated study are being deposited in in this Addgene (www.addgene.org/Guillaume_Jacquemet/). The mass spectrometry proteomics data have been deposited in the ProteomeXchange Consortium via the PRIDE (Perez-Riverol et al., 2022) partner repository with the dataset identifier PXD067430 (Project DOI: PXD067430). The raw microscopy images and the numerical data used to make the figures have been archived on Zenodo (https://zenodo.org/records/17779715). The authors declare that the data supporting the findings of this study are available within the article and from the authors upon request. Any additional information required to reanalyze the data reported in this paper is available from the corresponding authors.

## Manuscript preparation

Figures were prepared using Fiji and Inkscape. PlotsOfData (Postma and Goedhart, 2019). Volcano plots were generated with VolcaNoseR (Goedhart and Luijsterburg, 2020). SuperPlots were created using SuperPlotsOfData (Goedhart, 2021). GPT-5 (OpenAI) and Grammarly (Grammarly, Inc.) served as writing aids during manuscript preparation. The author also edited and validated all text sections. GPT-5 did not provide references. The PDF version of this manuscript was formatted with Rxiv-Maker (Saraiva et al., 2025).

## Competing interests

The authors declare that they have no competing or financial interests.

## Author contributions

**Conceptualization**: G.J. **Methodology**: A.P., N.J.B., M.M., O.J., M.D., M.O., G.J. **Reagents**: B.T.G., J.I. **Formal Analysis**: A.P., N.J.B., M.M., O.J., M.D., M.O., J.P., B.T.G., G.J. **Investigation**: A.P., N.J.B., M.M., O.J., M.D., M.O., J.P., B.T.G., G.J. **Writing – Original Draft**: A.P., G.J. **Writing – Review and Editing**: Everyone. **Visualization**: A.P., N.J.B., B.T.G., G.J. **Supervision**: B.T.G., G.J. **Funding Acquisition**: G.J.

## Acknowledgments

This study was funded by the Research Council of Finland (338537 and 371287 to G.J.), the Sigrid Juselius Foundation (to G.J. and J.I.), the Cancer Society of Finland (Syöpäjärjestöt; to G.J. and J.I.), and the Solutions for Health strategic funding for Åbo Akademi University (to G.J.). Additionally, this research received support from the InFLAMES Flagships Programme of the Research Council of Finland (decision numbers: 337530, 337531, 357910, and 35791) and Centre of Excellence (decision numbers: 346131 and 364182 to J.I.). J.I. is supported by the Finnish Cancer Institute (K. Albin Johansson Professorship). B.T.G. and N.J.B. received support from Cancer Research UK Program Grant (CRUK-A21671). O.J. was supported by the University of Turku Graduate School Doctoral Program in Technology (DPT) (2300668) and the Finland Fellowship 2023. M.D. received support from the European Union’s Horizon Europe research and innovation program under Marie Sklodowska-Curie grant agreement number 101108089. This work was supported by the Research Council of Finland, FIRI 2023 grant decision numbers 359073 and 358879, and FIRI 2024 grant decision numbers 367582 and 367577. Additionally, this project received funding from the European Research Council (ERC) under the European Union’s Horizon 2020 research and innovation program (Grant agreement No. 101142305 to J.I.). Testament funds from Henna Ruusunen also supported this work. Imaging was performed at the Cell Imaging and Cytometry Core at Turku Bioscience Centre, supported by the Finnish Advanced Microscopy Node of Euro-BioImaging Finland (Turku, Finland), and Turku Bioimaging. Mass spectrometry analyses were carried out at the Turku Proteomics Facility, which is supported by Biocenter Finland.

## ABOUT THIS MANUSCRIPT

This work is licensed under CC BY 4.0.

## Supplementary Information

**Sup. Fig. S1.**
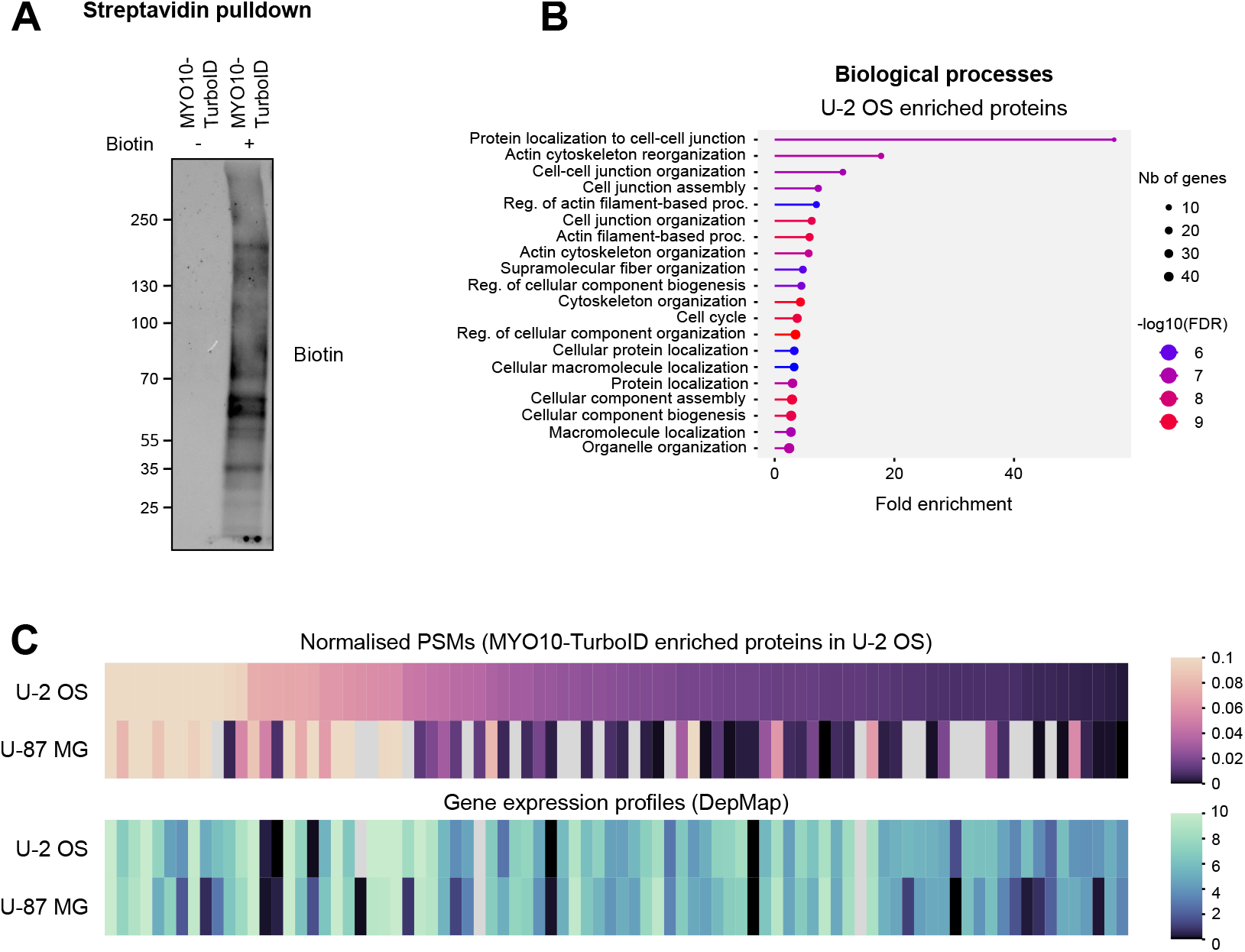
Identification of MYO10-associated proteins using proximity-dependent biotinylation - associated with Fig. 1. (**A**) U-2 OS cells stably expressing TurboID-MYO10 were cultured on fibronectin-coated dishes for 2 hours, with biotinylation initiated 1 hour before lysis. Biotinylated proteins were affinity-purified using streptavidin-coated beads and analyzed by western blotting. (**B**) Gene ontology analysis highlighting biological processes significantly enriched among proteins identified in U-2 OS cells. (**C**) Heatmaps comparing detected MYO10-associated proteins between the two cell lines, specifically highlighting those enriched in the U-2 OS dataset. Expected expression profiles derived from the DepMap database are included for comparison. The numerical data and the raw images used to make this figure have been archived on Zenodo (https://zenodo.org/records/17779715).

**Sup. Fig. S2.**
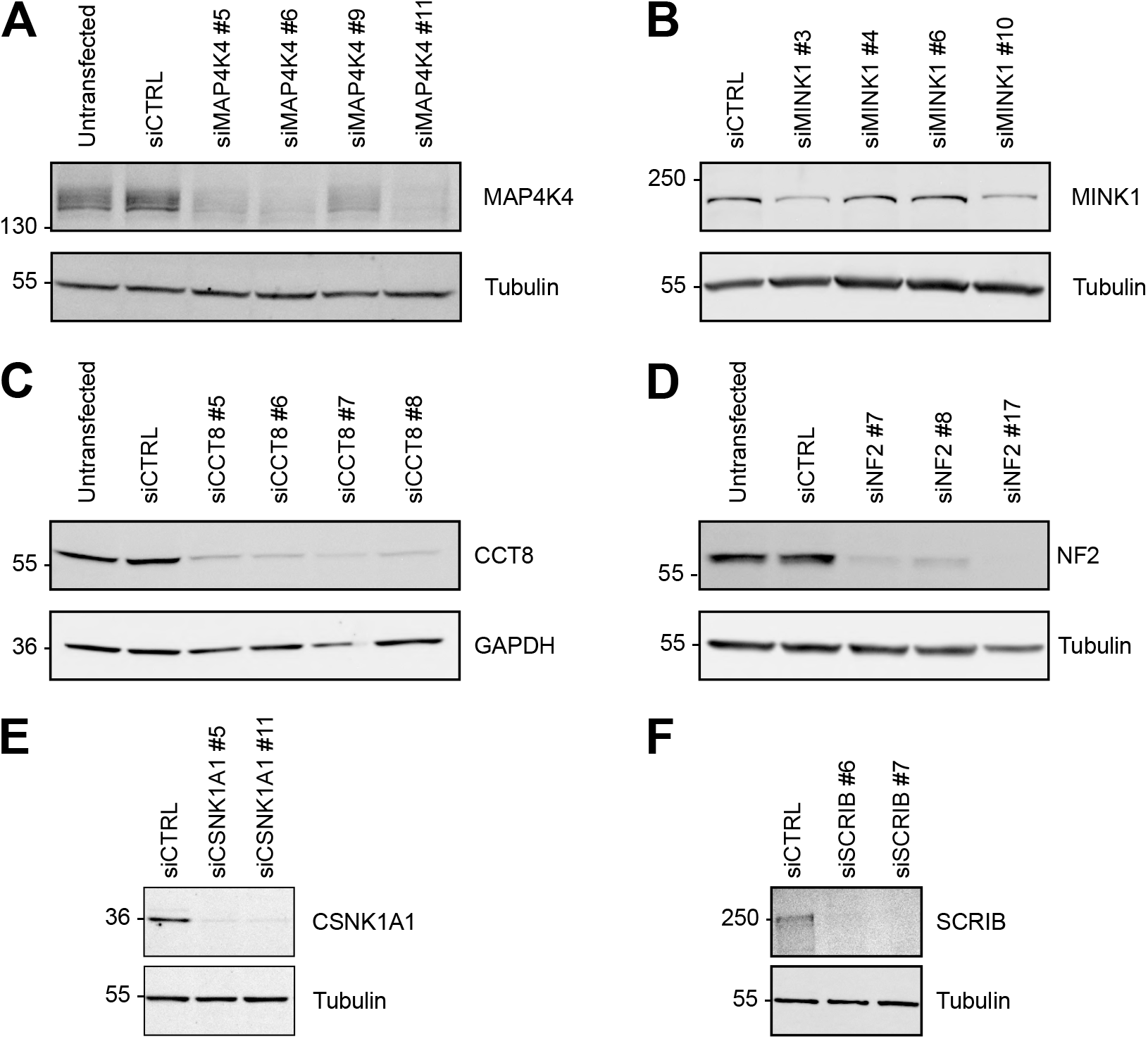
Validation of silencing efficiency using western blot - associated with Fig. 2. (**A**-**F**) U-87 MG cells were treated with multiple siRNAs targeting MAP4K4 (**A**), MINK1 (**B**), CCT8 (**C**), NF2 (**D**), CSNK1A1 (**E**), SCRIB (**F**), or a control siRNA (siCTRL). For each condition, cell lysates were collected, and siRNA efficiency was validated by western blot. Representative western blots are shown. The raw images used to make this figure have been archived on Zenodo (https://zenodo.org/records/17779715).

**Sup. Fig. S3.**
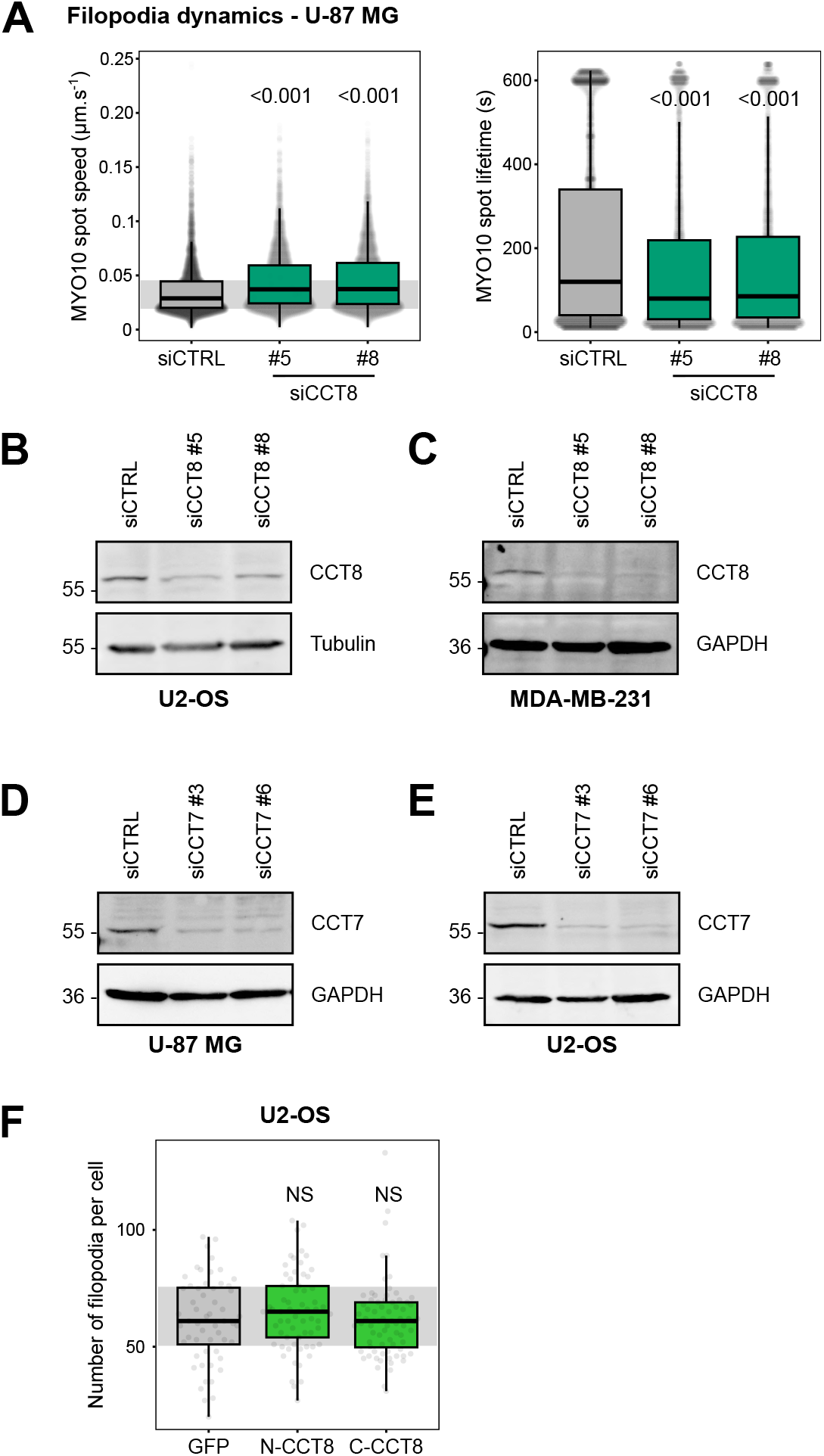
CCT8 modulates filopodia formation independently of the TRiC complex - associated with Fig. 3. (**A**) CCT8-silenced U-87 MG cells transiently expressing MYO10-GFP were plated on FN and imaged live using an Airyscan confocal microscope (1 picture every 5 s over 20 min). For each condition, MYO10-positive particles were automatically tracked. The average MYO10 track duration per cell is displayed (three biological repeats, n > 29 cells per condition). The average speed and the lifetime of MYO10 spots are shown (n > 5000 filopodia; three biological repeats). (**B, C**) U-2 OS (**B**) and MDA-MB-231 (**C**) cells were treated with siRNAs targeting CCT8 or a control siRNA (siCTRL). For each condition, cell lysates were collected, and siRNA efficiency was validated by western blot. Representative western blots are shown. (**D, E**) U-87 MG (**D**) and U-2 OS (**E**) cells were treated with siRNAs targeting CCT7 or siCTRL. For each condition, cell lysates were collected, and siRNA efficiency was validated by western blot. Representative western blots are shown. (**F**) Cells expressing mScarlet-MYO10 along with GFP alone, GFP-N-CCT8 (N-terminal tagged), or GFP-C-CCT8 (C-terminal tagged) were treated with siCTRL, plated on fibronectin-coated coverslips for 2 hours, fixed, and imaged using spinning disk confocal microscopy. Filopodia numbers per cell were quantified. (n > 65 cells per condition, three biological replicates). Results are shown as boxplots, with whiskers extending from the 10th to the 90th percentiles. The boxes illustrate the interquartile range, and a line inside each box indicates the median. Data points outside the whiskers are shown as individual dots. The numerical data and the raw images used to make this figure have been archived on Zenodo (https://zenodo.org/records/17779715).

**Sup. Fig. S4.**
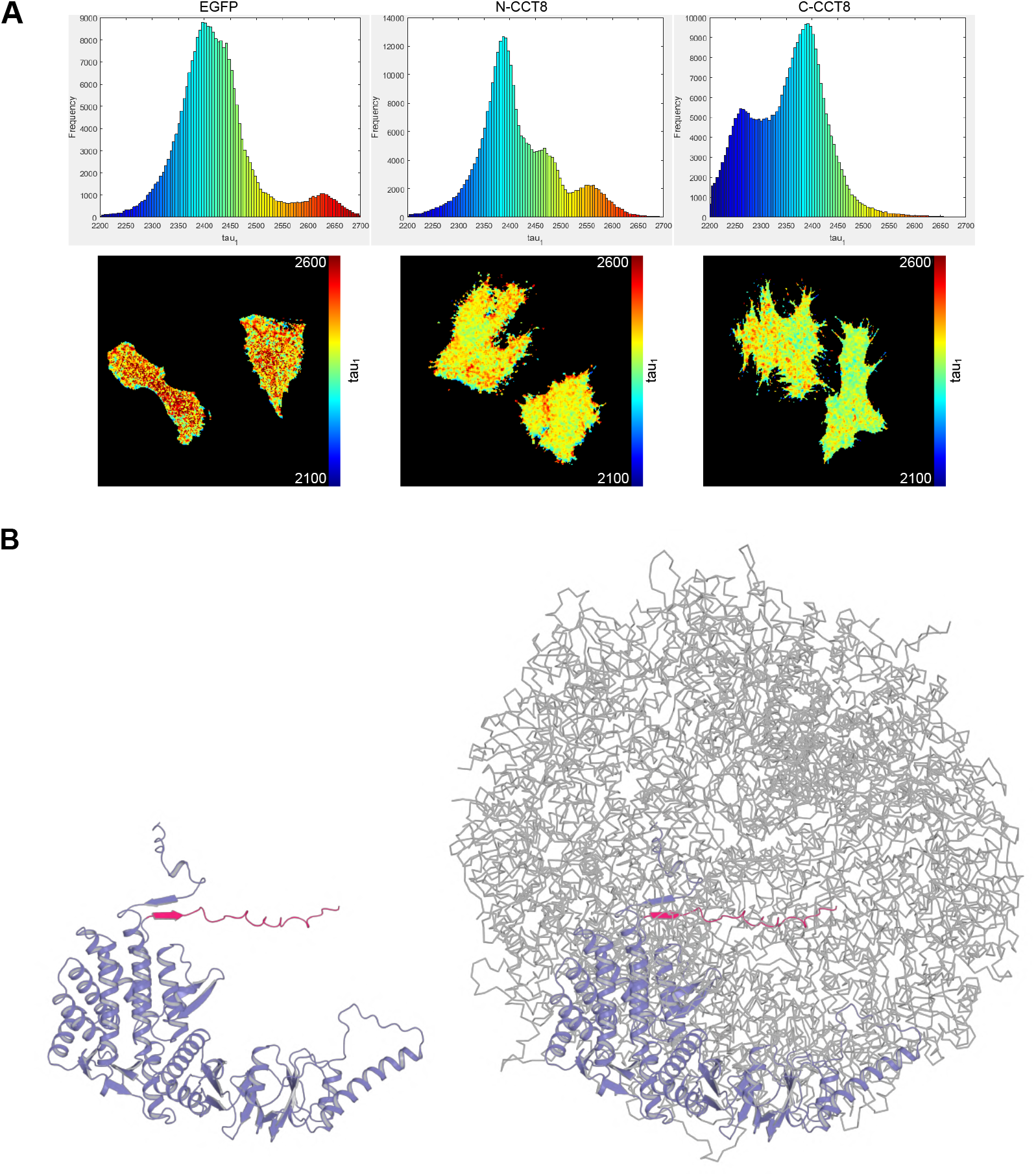
CCT8 binds to MYO10 - associated with Fig. 4. (**A**) GFP lifetime (τ) was measured in U-87 MG cells co-expressing MYO10–mScarlet with GFP, GFP–CCT8 (N-terminal tag), or CCT8–GFP (C-terminal tag). Representative images are displayed. (**B**) The C-terminus of CCT8 (residues 520-548; pink) is exposed (left) or buried (right), depending on whether CCT8 is part of the TRiC complex (6KS6, (Jin et al., 2019)). One copy of CCT8 is shown as a blue cartoon, and the other components of the TRiC complex are depicted as grey ribbons.

**Sup. Fig. S5.**
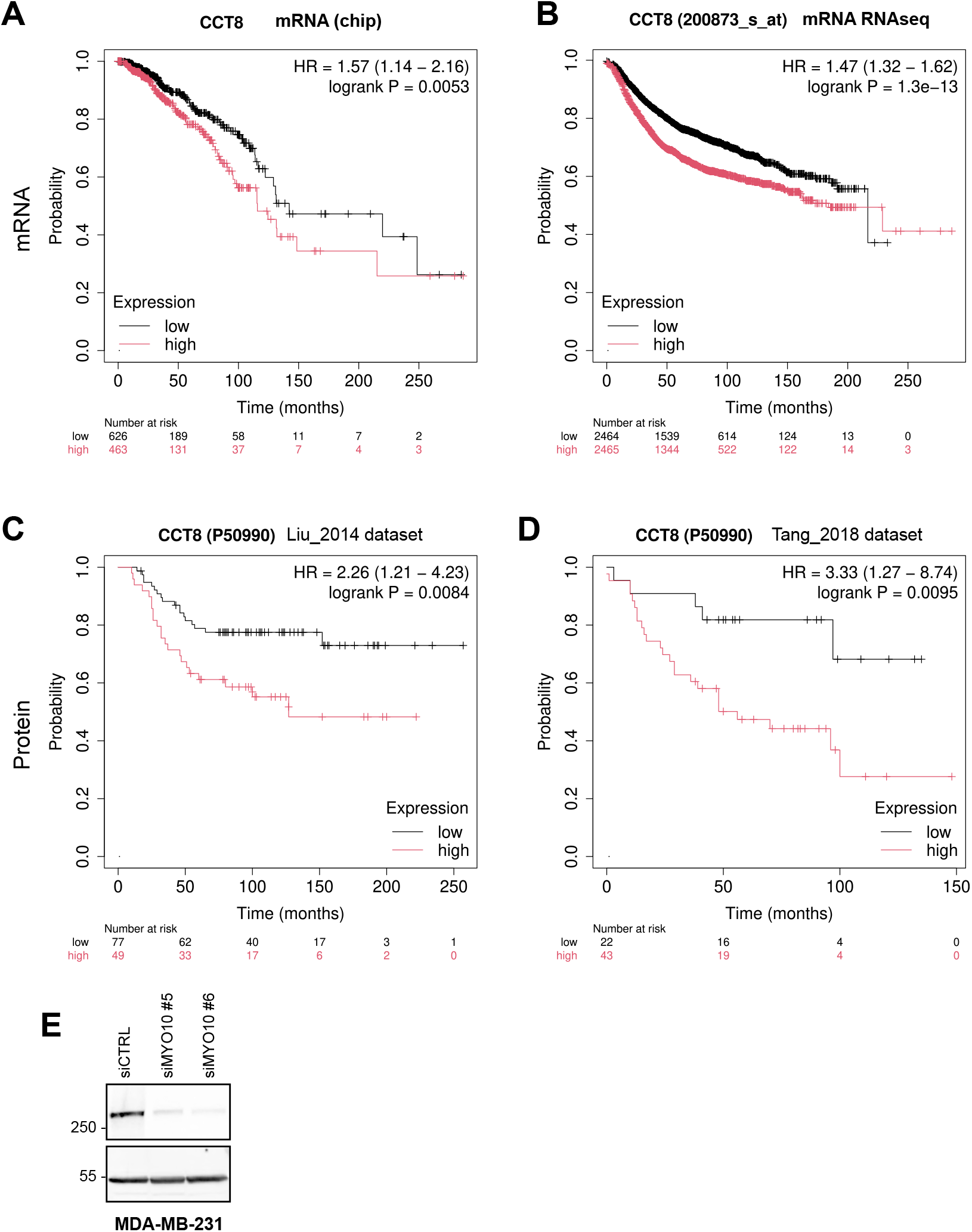
CCT8 expression and connection to prognosis in breast cancer patients and MYO10 silencing in MDA-MB-231 cells, associated with Fig. 5. (**A**-**D**) Kaplan-Meier survival analyses of breast cancer patients segregated into high or low CCT8 mRNA (**A** and **B**) or protein levels (**C** and **D**). These analyses were performed using the Kaplan-Meier plotter (Posta and Győrffy, 2025) and four different datasets, as indicated. (**E**) MDA-MB-231 cells were treated with siRNAs targeting MYO10 or with siCTRL. For each condition, cell lysates were collected, and siRNA efficiency was validated by western blot. A representative western blot is shown. The raw images used to make this figure have been archived on Zenodo (https://zenodo.org/records/17779715).

## Notes

https://zenodo.org/records/17779715

### Competing Interest Statement

The authors have declared no competing interest.

